# Structures and enzymatic mechanisms of DRT7/UG10 antiphage reverse transcriptases

**DOI:** 10.64898/2026.02.16.706125

**Authors:** Małgorzata Figiel, Vysakh Komathattu Viswanath, Mariusz Czarnocki-Cieciura, Markéta Šoltysová, Julia Rybakowska, Artyom A. Egorov, Vasili Hauryliuk, Marcus J. O. Johansson, Marcin Nowotny

## Abstract

The ongoing evolutionary arms race between bacteria and bacteriophages has driven the emergence of numerous anti-phage defense systems. Several of these defenses contain domains related to reverse transcriptases (RTs), DNA polymerases that utilize RNA as a template. Certain members of the Abi/UG RT family employ a unique enzymatic mechanism combining protein-primed DNA synthesis with template-independent polymerization. A subgroup, UG10 – also known as Defense-related RT 7 (DRT7) – is characterized by an enzymatic core comprising an RT-like domain and a primase domain, accompanied by a poorly characterized accessory element. This element is either C-terminally fused to the core or encoded by a separate open reading frame. Here, we present cryo-EM structures of protein–DNA conjugates from two UG10 enzymes, revealing a compact yet flexible domain architecture. We demonstrate that the poly(dT) product synthesized by the RT-like domain serves as a template for the primase domain to generate a complementary poly(A) strand. Combining biochemical experiments with structural modeling, we propose the conformational changes necessary for protein priming and coordination between the RT and primase domains. Finally, we show that DRT7 exhibits broad-spectrum anti-phage activity and demonstrate that the λ phage DNA mimic protein Gam can trigger this defense system.

## INTRODUCTION

Reverse transcriptases (RTs) are DNA polymerases uniquely characterized by their ability to use RNA as a template. Found in all domains of life, these proteins play key roles in the propagation of mobile genetic elements called retrotransposons^1^, viral proliferation^2^, and maintenance of telomeric ends in eukaryotic genomes^3^. Bacterial RTs are classified into several types^4^, including: 1) RTs associated with group II intron retroelements, 2) group II intron-like RTs, 3) RTs encoded in diversity generating retroelements^5^, 4) RTs associated with CRISPR-Cas systems^6^, 5) RTs encoded in Retron antiphage defenses^7,8^, 6) Abortive infection (Abi) antiphage defense RTs^9,10^, and, finally, 7) less explored and diverse Unknown Group (UG) RTs, which have recently been established as acting in antiphage defense as well^11–13^.

Abi and UG RTs together form a highly diverse Abi/UG clade^7^. In these proteins, the RT domain is either fused to additional domains or accompanied by noncoding RNAs (ncRNAs) or putative protein partners encoded by associated open reading frames (ORFs). The repertoire of fused or associated enzymatic domains is broad, encompassing nucleases, primases, methylases, and helicases. The functional role of these accessory elements remains poorly understood, but they may be coupled with RT activity to jointly produce a nucleic acid product or play regulatory roles, for example, by sensing phage infection.

The Abi/UG clade can be divided into four classes, further subdivided into 42 sub-groups^14^. Class 1 enzymes contain a C-terminally fused domain composed of α-helical repeats (αRep). This category includes all three known Abi RTs: AbiA, AbiK, and Abi-P2. Class 2 RTs lack fused domains but are accompanied by ncRNA or associated ORFs. Class 3 RTs are fused to or associated with nitrilase and phosphohydrolase domains. Remaining Abi/UG RTs fall into an unclassified category.

Recent studies have revealed a fascinating variety of unusual molecular mechanisms in class 2 UG enzymes, whose RT domains function as typical template-dependent reverse transcriptases^15,16^. DRT9, from the UG28 group, forms an oligomeric complex with its cognate ncRNA, which, in response to increased dATP levels, synthesizes poly(A) ssDNA proposed to sequester the ssDNA-binding protein of the phage^16–18^. In the DRT2/UG2 system, the RT domain utilizes its associated ncRNA as a template to generate a protein-coding complementary DNA (cDNA) with multiple repeats of the ncRNA sequence; subsequent translation of the ncRNA leads to cell growth arrest, halting phage propagation^15,19,20^. Tandem-repeat cDNAs are also produced by a UG17 system RT, DRT10, which has recently been proposed as the prokaryotic ancestor of telomerase^21^.

The first members of the Abi/UG RT family to be thoroughly characterized biochemically and structurally were Abi RTs^9,10,22^. Unlike most RTs, including class 2 of the UG clade, Abi enzymes do not utilize a template and can perform template-independent synthesis of single-stranded DNA (ssDNA)^9,10,22^. Furthermore, they do not require a nucleic acid primer; instead, they rely on protein priming, where the first nucleotide is attached to a tyrosine residue in the protein. While protein priming has also been described for hepatitis B virus RT^23^ and adenoviral DNA polymerase^24^, the combination with template-independent DNA polymerization is unique to Abi RTs. Technically, Abi RTs should be called RT-like enzymes, but for simplicity, we will refer to them as RTs throughout this work. The first structural insight into the workings of Abi/UG members came from structures of AbiK, Abi-P2, and AbiA antiphage defense systems^9,10^. A common feature of these structures is a bilobal, C-shaped arrangement of the catalytic domain and the C-terminal αRep domain. While divergent between families, αRep domains retain an overall solenoid topology. To be active, these RTs assemble into homo-oligomers, with different numbers of subunits: AbiA functions as a dimer, while AbiK and Abi-P2 form hexamers and trimers, respectively. Abi RTs possess a narrow central channel that accommodates only ssDNA. The AbiK-DNA structure visualized the binding of nascent DNA within this channel and its covalent attachment to a tyrosine residue, resulting from protein-primed synthesis^9^. Although conceptually similar, AbiK and AbiA differ in their biochemical properties, for example, exhibiting distinct nucleotide preferences^10^.

While originally proposed to act via altruistic suicide or growth arrest of the infected cell (abortive infection)^11–13^ recent studies suggest that these systems may directly interfere with phage replication rather than trigger cell death^10^. The antiphage defense function has been postulated for many UG RTs based on their predicted association with defense islands in bacterial genomes^14^, but experimental validation is lacking for most families^14^. UG RTs for which defense has been demonstrated have been renamed defense-associated reverse transcriptases (DRTs), numbered from type 1 to 11^14,21,25,26^. One such group is UG10, a member of Class 1 Abi/UG RTs, which has been named DRT7^14^.

In UG10/DRT7, the RT domain is fused with an N-terminal PriS primase domain, which belongs to the archaeo-eukaryotic primase superfamily^14^ (Fig. 1). A primase initiates nucleic acid synthesis on a template *de novo* without a primer. While a functional primase domain is required for the antiphage activity of DRT7 from *E. coli* SC366 strain against T2, T5 and ZL-19 phages^14^, its specific molecular function in defense remains unclear. The primase domain is connected to the RT domain via a primase large subunit C-terminal domain (PriL-CTD)^27^. The role of the primase domain remains enigmatic. Furthermore, members of the UG10 group carry an uncharacterized domain – either encoded in a separate ORF downstream of the RT gene (UG10 small or UG10S subfamily) or fused to the RT part at its C-terminus (UG10 large or UG10L subfamily) (Fig. 1).

**Figure 1.**
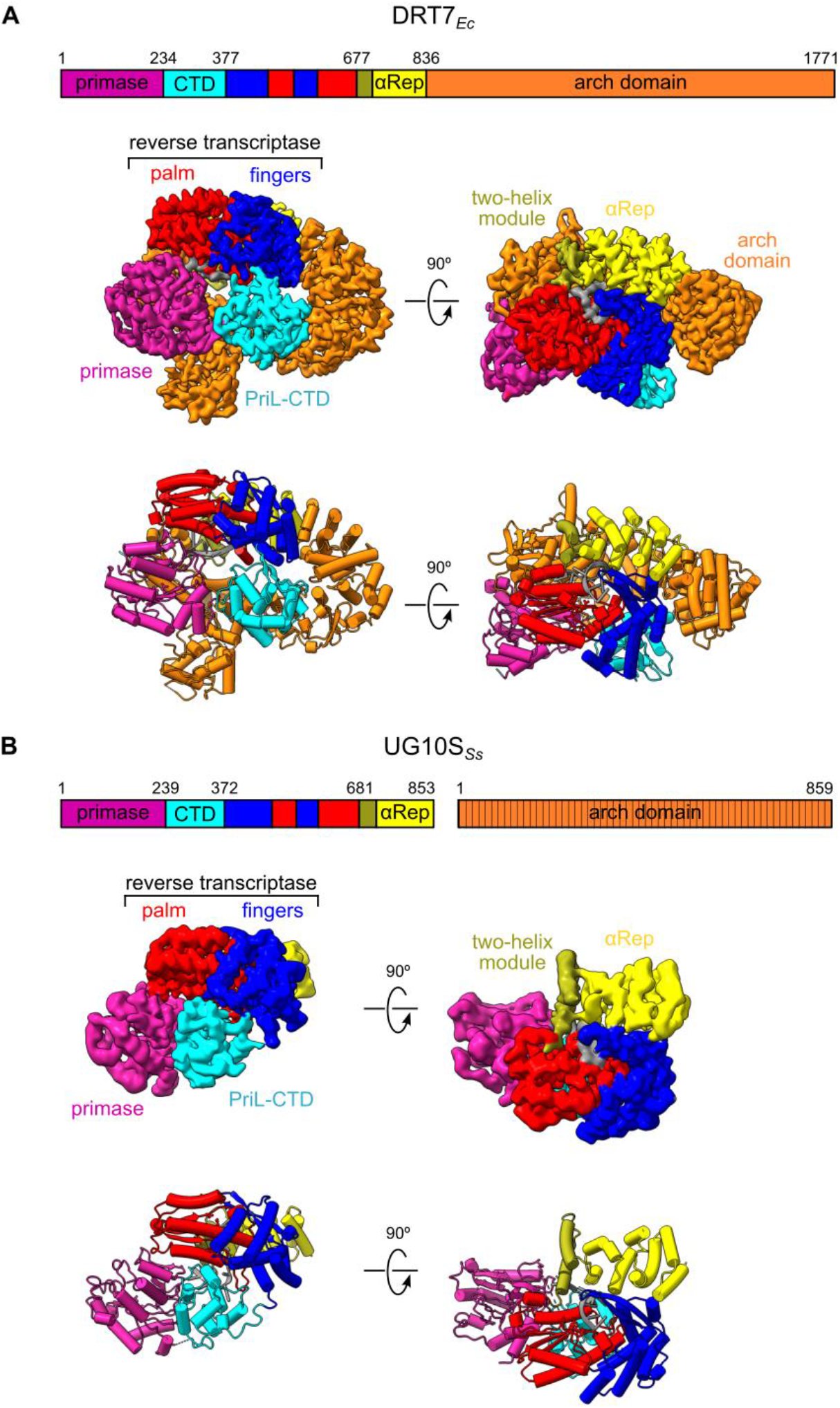
Overall structures of DRT7_*Ec*_ (A) and UG10S_*Ss*_(B). Schematic representations of domain composition are shown above cryo-EM reconstructions and corresponding structures (cartoon representation), with domains color-coded as indicated. Striped fill denotes the protein for which the structure could not be determined.

Here, we uncover the mechanism of UG10 RTs through structure-function studies of *E. coli* LN97 DRT7/UG10L (DRT7_*Ec*_) and *Streptococcus salivarius* UG10S (UG10S_*Ss*_). We present cryo-EM structures of DRT7_*Ec*_ and UG10S_*Ss*_ proteins and their variants, revealing the remarkable flexibility of these molecular machines and proposing a role for the previously uncharacterized C-terminal accessory domain, which we named the arch domain. We demonstrate that the RT and primase activities of the two UG10 RTs cooperate to produce double-stranded DNA (dsDNA) products, with the RT domain producing a poly(dT) strand used as a template by the primase domain to synthesize a complementary poly(dA) strand. Finally, we establish the defense profile of the DRT7_*Ec*_ system in *E. coli* and, through selection of DRT7_*Ec*_-insensitive escape mutants, show that the λ phage DNA mimic protein Gam can activate the defense system.

## RESULTS

### Comparative structural analysis of UG10 RTs reveals variations in the common architecture

Our first objective was to understand the molecular architecture of UG10 enzymes. We used DRT7 from *E. coli* strain LN97 (WP_247156398.1, hereafter DRT7_*Ec*_). The protein is readily purified upon overexpression in *E. coli*, and its sequence is nearly identical to previously characterized *E. coli* SC366 DRT7^14^, differing by only one amino acid (Gly instead of Asp in position 970). For comparative studies of UG10L vs. UG10S, we targeted *Streptococcus salivarius* UG10S (WP_104020715.1), which was previously identified by Mestre et al.^14^. Unfortunately, we were unable to produce its cognate accessory protein (WP_104020716.1) in soluble, monodisperse form, either in *E. coli* or insect cells. Furthermore, we could not construct a plasmid for P_*tet*_-driven expression of the UG10S-accessory protein operon for phage defense assays, and therefore, could not test its antiphage activity. Without confirmation of the defense function, we refrained from renaming the *S. salivarius* UG10 protein to a DRT and refer to this system as UG10S_Ss_.

We initially used mass photometry to study the oligomeric state of UG10S_*Ss*_ and DRT7_*Ec*_. The results showed that, at nanomolar concentration, both proteins were predominantly monomeric (Supplementary Fig. 1). Next, we determined cryo-EM structures of both proteins, achieving a 2.9 Å resolution reconstruction for DRT7_*Ec*_ and a 3.7 Å resolution reconstruction for UG10S_*Ss*_ (Fig. 1, Supplementary Figs. 2 and 3). Atomic models were built into these reconstructions (Fig. 1). The DRT7_*Ec*_ structure revealed well-resolved primase, PriL-CTD, RT, and αRep domains as well as the C-terminal accessory domain, with the exception of a region at its terminus comprising residues 1528-1681. The accessory C-terminal domain forms an arch-like structure—henceforth we termed it the arch domain (AD). A density for a continuous ten-nucleotide DNA strand, covalently attached to the protein, was also clearly defined. For UG10S_*Ss*_, the entire polypeptide chain (primase, PriL-CTD, RT, and αRep domains) was resolved except for the loop connecting the primase and PriL-CTD domains (residues 237-247). Density was also observed for ten nucleotides of bound ssDNA, nine of which could be reliably modeled.

The cores of UG10S_*Ss*_ and DRT7_*Ec*_ structures comprise a primase domain followed by a small, globular PriL-CTD domain, an RT domain, and an αRep domain. The RT domain exhibits the canonical right-hand fold, with palm and fingers subdomains, but lacks the thumb subdomain, as previously observed for other Class 1 UG RTs^9,10,28,29^. The enzymatic cores of UG10S_*Ss*_ and DRT7_*Ec*_ are similar; superposition of individual domains yields rmsd values of 1.2–2.0 Å for the primase, PriL-CTD, and RT domains. The αRep domains differ more substantially, with an rmsd of 5.9 Å over 144 Cα atoms. Compared to UG10S_*Ss*_, the palm subdomain of DRT7_*Ec*_ contains an additional helix (residues 486-491) positioned next to the first helix of its fingers subdomain. Within the fingers subdomain, the third helix (residues 416-428 in DRT7_*Ec*_ and 411-426 in UG10S_*Ss*_) adopts a different position in the two structures, and the fifth helix in UG10S_*Ss*_ (residues 541-556) is replaced in DRT7_*Ec*_ by two shorter helices positioned roughly perpendicularly. The αRep domains also show minor differences in the arrangement of several helices; notably, the first two helices, termed the two-helix module, exhibit a shift in position, the significance of which is discussed below.

The RT domains of UG10S_*Ss*_ and DRT7_*Ec*_ are similar to those of founding members of Class 1 Abi/UG RTs – Abi proteins, including the well-characterized *Lactococcus lactis* AbiK (*Ll*-AbiK) (Supplementary Fig. 4). Superimposing the RT core of DRT7_*Ec*_ (palm and fingers subdomains) onto the corresponding regions of AbiK RT yields an rmsd of 4.4 Å (218 Cα atoms). The αRep domains of UG10 and AbiK are both α-helical and possess similar positions relative to the RT domains. However, the two structures differ significantly and are not superimposable (Supplementary Fig. 4).

### The structures reveal the molecular architecture of primase, PriL-CTD and arch domains

The closest structural homologue of DRT7_*Ec*_/UG10S_*Ss*_ identified by DALI^30^ was the CRISPR-associated Prim-Pol from *Marinitoga* sp. 1137. One of its structures corresponds to a complex with dsDNA product (PDB ID: 7NQF)^31^. Superimposing the CRISPR-associated Prim-Pol primase domain onto the corresponding domains of UG10 proteins yielded rmsd values of 1.6 Å over 168 Cα atoms for DRT7_*Ec*_ primase domain and 1.9 Å over 172 Cα atoms for UG10S_*Ss*_, indicating a high degree of similarity.

The primase domain is followed by a PriL-CTD domain, composed of α-helices and two short anti-parallel β-strands originating from the N- and C-termini of the domain. PriL-CTD is connected to neighboring domains through flexible linkers. The structure of this domain exhibits appreciable similarity to C-terminal domains of large primase subunits. DALI searches identified the respective regions in *S. cerevisiae* large primase (PDB ID: 8FOC, rmsd of 2.5 Å over 68 Cα atoms)^32^ and the polymerase alpha-primase complex of *X. laevis* (PDB ID: 8G99, rmsd of 3.5 Å over 85 Cα atoms)^33^.

Despite the high similarity of the individual domains, superimposing the entire primase-PriL-CTD-RT-αRep regions of DRT7_*Ec*_ and UG10S_*Ss*_ reveals larger differences. This is due to the distinct positioning of the primase and PriL-CTD domains relative to the RT domain, facilitated by flexible linkers connecting these three domains. In the UG10S_*Ss*_ structure, the PriL-CTD is located near the interface between the palm and fingers subdomains of the RT, forming tight interactions with both (Supplementary Fig. 5A). This positions the primase domain close to the palm subdomain and in the vicinity of the two-helical module of the αRep (Supplementary Fig. 5B). Conversely, in DRT7_*Ec*_, the primase, PriL-CTD, fingers, and palm subdomains are arranged in a square plan, with PriL-CTD positioned opposite the fingers subdomain and primase opposite the palm subdomain (Supplementary Fig. 5A). Moreover, the primase domain is farther from the αRep domain, resulting in a more open arrangement (Supplementary Fig. 5B). Consequently, the distance between the primase and the two-helical module of the αRep increases substantially. The interdomain interfaces of these regions are detailed in Supplementary Text.

In the DRT7_*Ec*_ structure, we visualized most of the C-terminal arch domain (Fig. 1A, Supplementary Fig. 6A). Five globular modules/subdomains can be identified (Supplementary Fig. 6A,B), described in greater detail in the Supplementary Information. Module 1 is a continuation of the αRep solenoid structure. Modules 2 and 4 interact with the PriL-CTD and primase domains, respectively. Module 3, spanning residues 1272-1468, is the only AD module that contacts several domains from the core of DRT7_*Ec*_. It interacts with the RT domain and the region around the primase active site, introducing a steric obstacle between the primase and the two-helix module of αRep (Supplementary Fig. 6E,F). In this conformation, the AD likely blocks potential cooperation between the enzymatic activities of the primase and RT domains (Supplementary Fig. 6D). The region spanning residues 1528-1681 was not defined in the cryo-EM map, but AlphaFold3 predictions suggest it forms a compact domain (module 5) protruding from module 4 of the AD (Supplementary Fig. 6B).

### The protein-primed RT polymerase domain of UG10 enzymes cooperates with the primase domain to synthesize dsDNA

To study the biochemical properties of UG10S_Ss_ and DRT7_*Ec*_, we performed a series of enzymatic assays. Recombinant DRT7_*Ec*_ and UG10S_Ss_ preparations had 260/280 nm absorbance ratios higher than the expected 0.6 for nucleic acid-free proteins – 0.71-0.73 for UG10S_*Ss*_ and 0.65-0.71 for DRT7_*Ec*_. This indicates that, despite nuclease treatment, the proteins co-purify with nucleic acid, suggesting that UG10 RTs, like other characterized members of Class 1 Abi/UG enzymes^9,10,28^, utilize protein priming resulting in the covalent attachment of the nascent DNA to the RT. This covalently attached DNA was also visualized in our cryo-EM structures.

We first tested the polymerase activity of wild-type (wt) UG10S_Ss_ and a primase-inactivated variant with two substitutions in the primase active site (D72A and D74A – hereafter referred to as AXA). The proteins were supplemented with individual dNTPs, and reaction products were resolved on a TBE-urea gel and stained with SYBR Gold nucleic acid dye (Fig. 2A). Wt proteins generated ~100 nt products with dTTP, whereas dATP yielded larger (~700 nt) and more abundant products. No products were observed with dCTP or dGTP. The AXA variant produced only a ~100 nt product with dTTP. We interpret these results as evidence that RT, the only active polymerase in the AXA variant, is a poly(dT) polymerase. However, wt protein also generated a poly(dA) product. Since only the wt protein contains an active primase domain, we propose that this additional product results from primase activity using a short poly(T) fragment co-purified with UG10S as a template. The longer poly(dA) products may arise from template slippage. This interpretation is supported by a recent preprint demonstrating that UG10 can produce partially double-stranded nucleic acid primarily composed of [poly(dT)/poly(dA)]^27^.

To enhance our experimental readout, we established a fluorescence-based RT activity assay using UG10S_*Ss*_ and DRT7_*Ec*_ (Fig. 2B). Proteins were mixed with substoichiometric amounts of fluorescein-dUTP, facilitating attachment of a single fluorescent dUMP molecule to the short DNA product. Then, an excess of dTTP was added to allow further extension. Samples were collected throughout a 60-minute time course, treated with EDTA and proteinase K, resolved on a TBE-urea gel, and visualized for fluorescence. Incubation with dTTP resulted in a lengthening smear of DNA fragments. UG10S_*Ss*_ was more active than DRT7_*Ec*_, but it generated shorter products (approx. 100 and 200 nt, respectively) (Fig. 2C).

**Figure 2.**
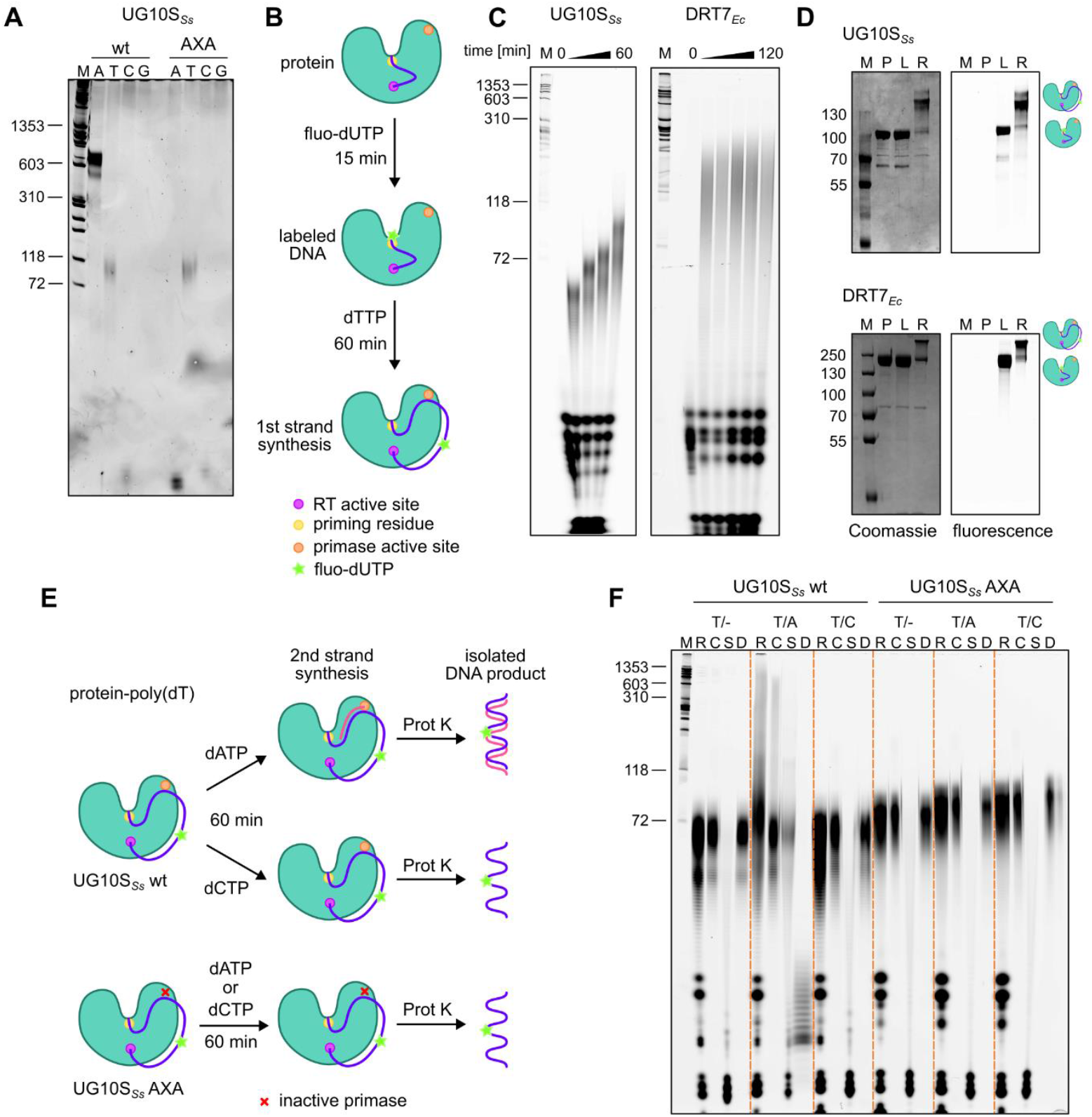
RT and primase activities of UG10 RTs cooperate to synthesize dsDNA. **A**) Denaturing TBE-urea polyacrylamide gel electrophoresis of reaction products from UG10S_*Ss*_ and its primase-deficient AXA variant, using individual deoxynucleotides and stained with SYBR Gold. (**B**) Schematic of a fluorescent dUTP-based DNA synthesis assay, illustrating the initial protein-primed DNA fragment and its labeling with fluorescein-coupled dUTP. UG10 co-purified with a short DNA fragment (blue line) attached to the protein priming residue (magenta circle) is incubated with fluorescein-coupled dUTP (green star) for labeling. Next, the DNA strand is extended in a reaction with dTTP. Polymerase and primase active sites are shown as yellow and orange circles, respectively. (**C**) Time course of polymerase activity of UG10 proteins using the assay in (B). (**D**) Visualization of covalent nucleic acid product attachment to UG10 RTs. Proteins were first labeled with fluorescein-dUTP and then incubated with an excess of dTTP for 1 hour. Samples were analyzed by SDS-PAGE and Coomassie blue staining (protein) and fluorescence detection (nucleic acid). M – size marker, P – protein, L – labeled sample, R – reaction sample. (**E**) Schematic of the dsDNA synthesis assay. The reaction starts with a protein-poly(dT) complex produced as depicted in (B). The complex is incubated with dATP or dCTP. In the presence of dATP, synthesis of complementary strand (pink line) is predicted to occur. Synthesis of the second strand is not expected for wild-type protein in the presence of dCTP nor in any reactions for the primase-inactivated variant. (**F**) Analysis of the products of UG10S_*Ss*_ in fluorescence-based assay depicted in (E). The nucleic acids were analyzed on a TBE-urea gel scanned for fluorescence. The proteins used (wt and AXA variant) are given on top of the gel. Nucleotide added [dTTP (T), dTTP and dATP (T/A) or dTTP and dCTP (T/C)] are indicated on top of each group of lanes. M – size marker, R – reaction product, C – purified reaction product, S – product treated with ssDNA-specific S1 nuclease, D – product treated with dsDNase.

We further analyzed samples from reactions with fluorescein-dUTP and dTTP by denaturing SDS-PAGE. Protein bands were visualized by both Coomassie blue staining and fluorescent signal, indicating covalent dUTP attachment to the protein. Following incubation with dTTP, protein bands exhibited a slower mobility – likely due to the covalent attachment of a longer poly(dT) DNA chain (Fig. 2D).

We then used the fluorescence-based assay to investigate whether the primase and RT domains cooperate to synthesize dsDNA. Unfortunately, difficulties preparing an expression construct for a primase-deficient DRT7_*Ec*_ variant prevented its use. We therefore focused on UG10S_*Ss*_, for which we generated the primase-inactive AXA variant as a control. The assay was performed as described above (Fig. 2B), with the addition of dATP or dCTP to the reaction (Fig. 2E). We hypothesized that primase would only utilize dATP when using poly(dT) as a template. Incubation with dATP resulted in a slightly different appearance of the DNA product on the gel, likely due to incomplete denaturation of poly(dT)/poly(dA) products (Fig. 2E). We next purified and analyzed the composition of reaction products using ssDNA- and dsDNA-specific nucleases – S1 nuclease and dsDNase, respectively (Fig. 2F). S1 nuclease digested the products of all reactions for both wt UG10S_*Ss*_ and the AXA variant. Degradation was less efficient for wt UG10S_*Ss*_ incubated with dTTP and then dATP. DsDNase could only digest the product from wt UG10S_*Ss*_ incubated with dTTP and dATP, confirming the presence of dsDNA. This product was also partially digested by S1 nuclease, suggesting that the complementary strand may have been synthesized as shorter fragments separated by single-stranded gaps. These results further support the idea that the primase and RT domains collaborate to produce dsDNA.

### Binding of the DNA product by DRT7_*Ec*_ and UG10S_*Ss*_

In both DRT7_*Ec*_ and UG10S_*Ss*_ structures, we observed additional density corresponding to a 10-nt ssDNA stretch co-purified with the protein (Fig. 3A). In the UG10S_Ss_ structure, the nucleotide in position 3 could not be modeled due to poor map quality in that region. Continuous DNA density in EM potential maps was connected to the hydroxyl groups of Tyr682 (DRT7_*Ec*_) and Tyr691 (UG10S_*Ss*_) (Fig. 3B), demonstrating a covalent linkage with the 5′-phosphate of the DNA and involvement in protein priming. Both residues are located on the first α-helix of αRep, within the two-helix module.

**Figure 3.**
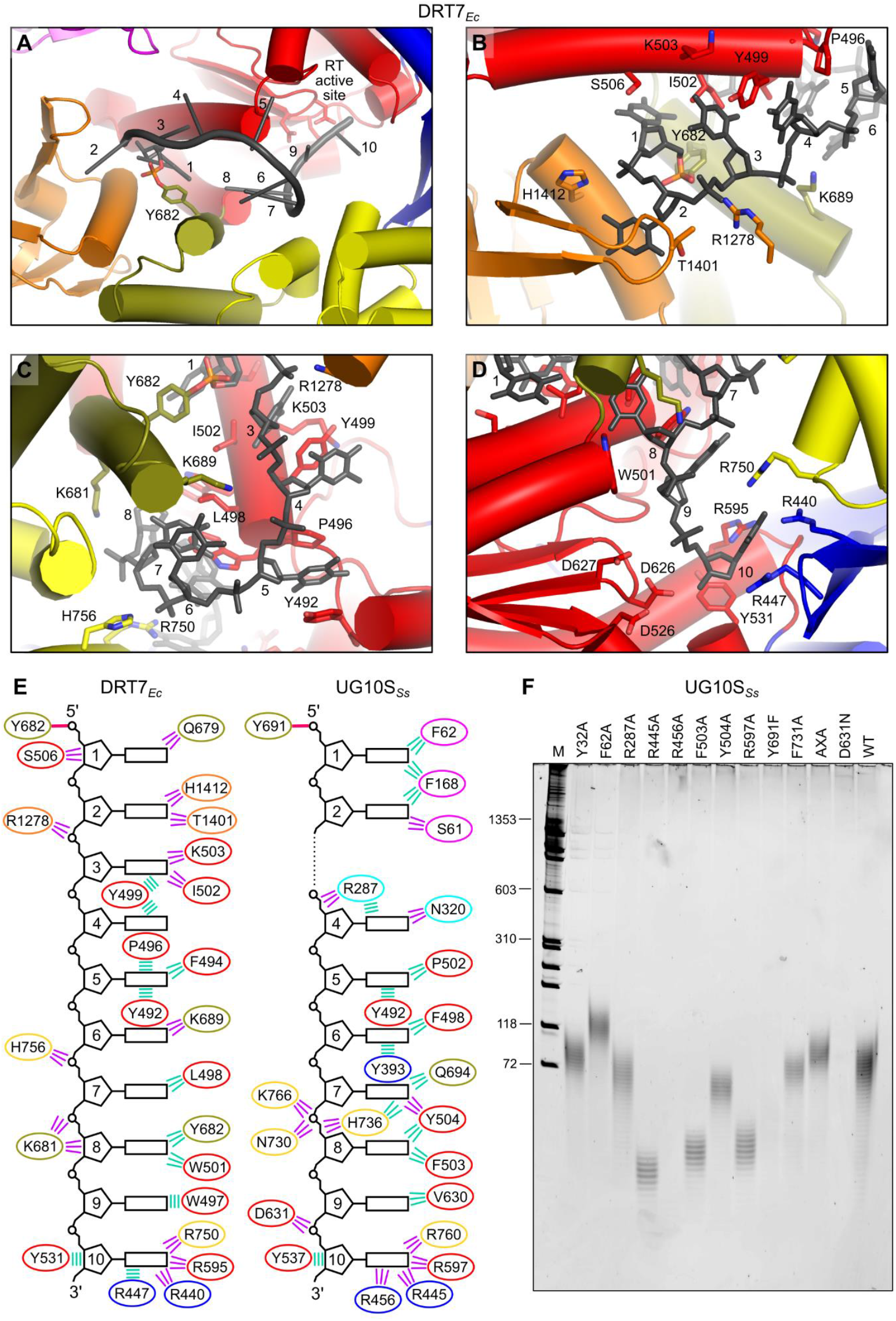
Protein-DNA contacts in UG10 proteins. (**A**) Overall view of ssDNA bound to DRT7_*Ec*_. ssDNA is shown as a black ladder; protein domains are colored as in Fig. 1. The priming tyrosine and RT active site residues are shown as sticks and labeled. (**B-D**) Close-up views of interactions between ssDNA and protein residues. DNA and interacting residues are shown as sticks. (**E**) Schematic of protein-DNA interactions observed in the structures. Ovals represent interacting protein residues (color-coded by domain as in Fig. 1). Thick pink lines indicate covalent linkages between priming tyrosines and the 5′ ends of the DNA products. Polar and van der Waals interactions are indicated by violet and green lines, respectively. (**F**) Polymerase activity of selected UG10S_*Ss*_ monitored by TBE-urea polyacrylamide gel electrophoresis with SYBR Gold staining.

All visible nucleotides form contacts with protein residues, with interactions mediated primarily by nucleobases. We number the nucleotides starting with the 5′ nucleotide attached to the priming tyrosine residue (nt 1), using DRT7_*Ec*_ residue numbering unless otherwise stated. The base of nt 2 is perpendicular to that of nt 1, creating a sharp turn in the DNA trajectory (Fig. 3B). The DNA then adopts a straight trajectory until nt 5, where the backbone makes another turn (Fig. 3C). The conformations of nucleotides 4 and 5 are stabilized by numerous contacts, including stacking interactions between nucleobases and aromatic sidechains, such as the conserved tyrosine in the palm subdomain (position 492) or proline 496 (DRT7_*Ec*_) (Fig. 3C). From nt 6, the DNA strand runs between the RT and αRep domains toward the RT polymerase active site. Numerous protein-DNA contacts, mainly through nucleobases, are formed by nts 7 and 8. Sidechains of αRep residues also interact with the DNA backbone flanking nt 7 (Fig. 3C). The base of nt 9 forms a stacking interaction with Trp497 (DRT7_*Ec*_) or a van der Waals contact with Val630 (UG10S_*Ss*_). The last modeled nucleotide is positioned at the RT active site, representing a pre-translocation state (Fig. 3D). Several conserved arginine sidechains contact this nucleotide: Arg750 (αRep), Arg595 (palm subdomain), and Arg440 and Arg447 (fingers subdomain) (Fig. 3D). The guanidinium group of Arg447 forms a π-π interaction with the nucleobase of nt 10. Additionally, a conserved tyrosine residue (Tyr531) interacts with the deoxyribose ring of nt 10. This contact would be much less efficient for a ribose ring comprising a 2′OH group, suggesting that Tyr532 acts as a ‘steric gate’ conferring specificity for deoxyribonucleotides and making DRT7/UG10 RTs DNA polymerases. The protein-DNA interactions observed in the structures of DRT7_*Ec*_ and UG10S_*Ss*_ are schematically shown in Figure 3E.

We then investigated the importance of residues involved in DNA binding, using UG10S_*Ss*_ as a model system. We generated protein variants with point substitutions of residues making key DNA contacts, along with the RT-inactive D631N variant and the primase-inactive AXA variant as controls. Polymerization activity was tested using an assay where dTTP was added to the protein, and resulting DNA products were analyzed on SYBR Gold-stained TBE-urea gels (Fig. 3F). As expected, the RT-inactive variant showed no activity, and the AXA variant displayed activity comparable to wt protein. Substitution of Tyr691 with phenylalanine completely abolished DNA synthesis, confirming the essential role of its hydroxyl group in protein priming. All other substitutions affected synthesis efficiency, impacting the amount and/or length of the ssDNA products. The magnitude of the effect correlated with the nucleotide position of the interacting residue, with the strongest effects observed at the 3′ end of the strand and near the RT active site. Complete abrogation of activity was observed for R456A. Arg456, like its DRT7_*Ec*_ counterpart Arg447, interacts with the incoming nucleotide and stabilizes its position through a stacking interaction at the active site. Alanine substitutions of the other arginine residues contacting the 3′-terminal nucleotide (Arg445 and Arg597) resulted in shorter products, as did the F503A substitution (UG10S_*Ss*_, interacts with nt 8 base). Limited effects on efficiency or length were observed with alanine substitutions of Tyr504 and Phe731, which form stacking interactions with the base of nt 7. A similar, but slightly more pronounced, effect on DNA production was observed for the R287A variant. The guanidinium group of Arg287 stacks against the nucleobase at position 4 and is one of the two residues of the PriL-CTD domain involved in contacts with the DNA product of the RT domain.

Finally, we examined the role of two residues in the primase domain involved in template DNA binding. Substitution variants were created for Phe62, which engages in a stacking interaction with the first nucleotide attached to the priming residue, and Tyr32, a conserved residue critical for the initiation stage of primase-catalyzed DNA synthesis. The amount of DNA produced by the Y32A variant was slightly decreased. The product of the F62A variant was longer than for the wild-type, suggesting that the absence of the stacking interaction with the 5′ end of the DNA strand conferred conformational flexibility, facilitating extension.

### Structural rearrangements involved in protein priming and primase activity

The priming tyrosines of UG10 proteins are located within the two-helix modules of the αRep domains. For priming (i.e., covalent attachment of the first nucleotide) to occur, these residues need to be positioned at the RT active site, which is likely facilitated by the flexibility of the two-helix module. To capture the conformation preceding DNA synthesis, we analyzed a cryo-EM structure of an RT-inactive UG10S_*Ss*_ variant (D631N). Consistent with the absence of attached nucleic acid, the 260/280 nm absorbance ratio of this protein was 0.5 (Supplementary Fig. 7). Cryo-EM reconstructions revealed a well-defined primase-RT portion but lower-resolution maps of the αRep region, indicative of substantial conformational flexibility (Fig. 4A, B). This suggests that covalent attachment of DNA stabilizes the αRep domain’s position.

**Figure 4.**
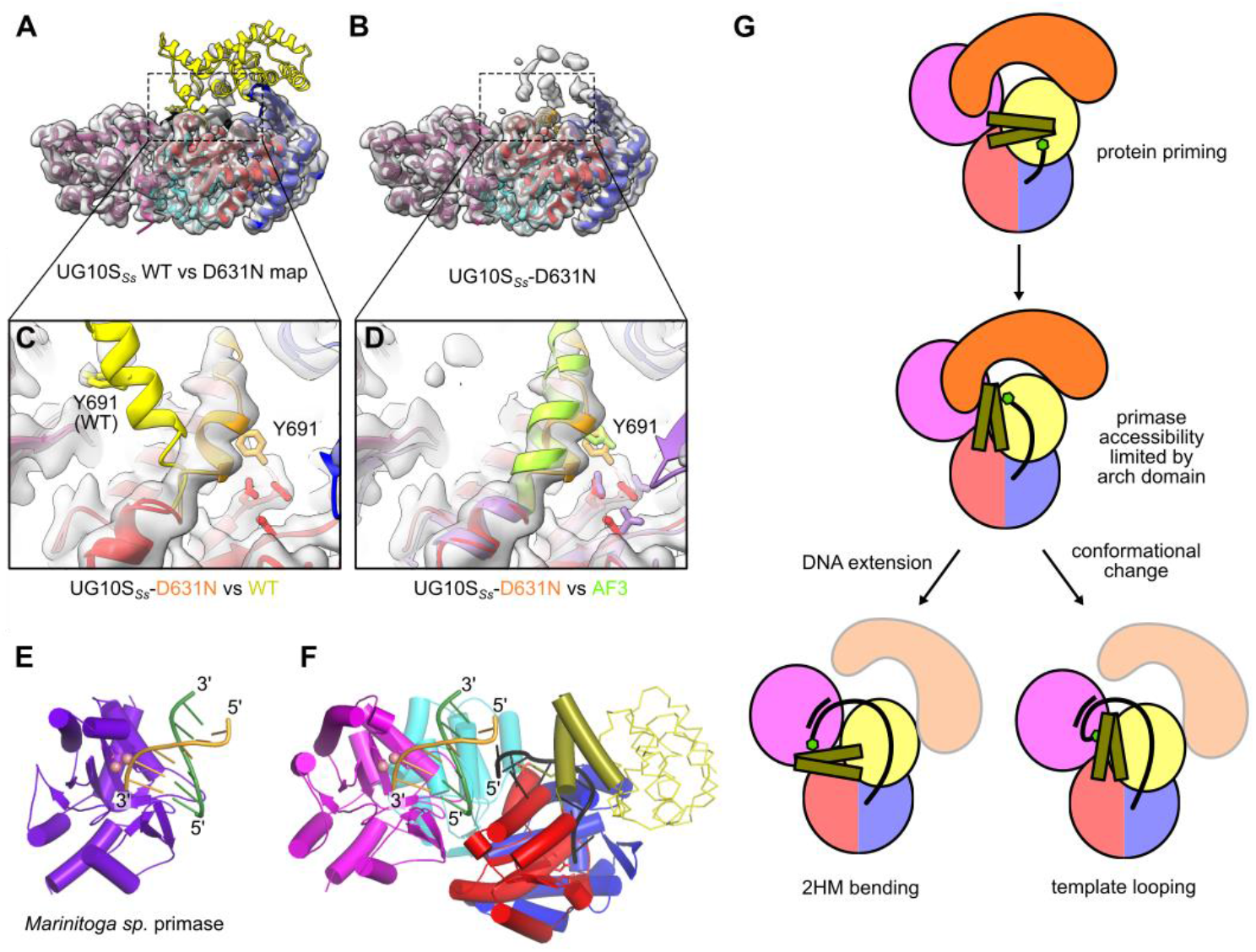
Conformational changes enabling protein-priming and primase activity of UG10S_*Ss*_. (**A, B**) Cryo-EM map of the polymerase-deficient UG10S_*Ss*_ D631N variant overlaid on the wild-type protein structure (A) and the D631N structure (**B**). Domains are color-coded as in Fig. 1. (**C, D**) Close-up views of the protein-priming helix region. UG10S_*Ss*_ D631N structure is superposed on the wild-type structure (C) and on an AlphaFold3 model (D). The protein-priming helix of UG10S_*Ss*_ D631N is shown in orange; the AlphaFold model is shown in violet and green. (**E**) Structure of a bacterial primase from *Marinitoga* sp. complexed with double-stranded DNA (PDB ID: 7NQF^31^). Template and product strands are shown in green and orange, respectively. Active site residues and coordinated cobalt ions are shown as sticks and pink spheres, respectively. (**F**) Model of dsDNA synthesis by UG10 enzymes. The primase domain is oriented as in (E); DNA and cobalt ion positions were predicted by superimposing 7NQF onto the UG10S_*Ss*_ primase domain. The two-helix module is shown in cartoon representation, and the remaining αRep domain is shown in wire representation. The template strand is shown in green. (**G**) Schematic of conformational rearrangements required for protein-priming and primase activity. RT-initiated DNA synthesis requires repositioning of the two-helix module. Access to the primase domain appears to be obstructed by the arch domain, which must be displaced for complementary strand synthesis. The ssDNA product may be positioned for primase activity through bending of the two-helix module or looping of the elongated DNA strand.

Notably, the UG10S_*Ss*_-D631N reconstruction exhibited density not present in the wild-type structure: an extension of a short helical region at the end of the palm subdomain. The shape of this density corresponded to an α-helix running approximately parallel to the 503-519 helix of the palm subdomain, directly above the β-sheet harboring active site residues. We modeled residues 678-695 from the first αRep helix, containing the protein-priming tyrosine, into this density (Fig. 4C, D).

Curiously, the hydroxyl group of this tyrosine residue is located at the RT active site near the position which would be required for the attachment of the first DNA nucleotide. Additional stabilization of the tyrosine is likely through interactions with the aromatic sidechains of neighboring residues Trp507 and Tyr629. Interestingly, one of the 25 models generated by AlphaFold3 also showed this conformation (Fig. 4D), further supporting its relevance. These analyses demonstrate that the αRep domain exhibits substantial conformational flexibility, and attachment of the first nucleotide to the priming tyrosine requires rearrangement of the two-helix module relative to the rest of the structure.

We then investigated how the UG10 primase domain might synthesize dsDNA. Using the structure of *Marinitoga* sp. primase complexed with dsDNA (Fig. 4E), we modeled the trajectory of the ssDNA product from the RT domain to a position where it could serve as a template for complementary strand synthesis by the primase (Fig. 4F). Assuming that the 3′ end of the ssDNA remains bound near the RT active site, we predict two possible scenarios. In our structures, the two-helix priming module can adopt different positions. Its further repositioning towards the primase domain could bring the DNA covalently attached to the priming tyrosine close to the primase active site (Fig. 4G). Alternatively, an appropriate DNA trajectory could be achieved without additional conformational changes by looping out the elongated ssDNA. However, for DRT7_*Ec*_, both scenarios would require displacement of the arch domain (AD) (Fig. 4G). In the DRT7_*Ec*_ structure, the tip of the AD is positioned between the primase domain and the 5′ portion of the DNA, blocking access to the primase active site. It also prevents movement of the two-helix module to present the ssDNA to the primase. This suggests that the AD regulates primase activity.

In summary, protein priming of the first strand involves a conformational rearrangement of the two-helix module containing the priming tyrosine. We also predict that further rearrangements are required for the primase to synthesize the complementary strand. Overall, this underscores the importance of flexibility for the UG10/DRT7 molecular machinery.

### DRT7_*Ec*_is a broad-spectrum antiphage defense system that can be activated by the phage DNA mimic protein Gam

DRT7 from *E. coli* strain SC366 has demonstrated defense activity against phages T2, T5 and ZL-19^14^. To further characterize this activity, we tested DRT7Ec against a broader range of phages. The *drt7* gene was cloned under the constitutive P_*tet*_ promoter in a pBR322-derived vector^34^, and liquid growth assays showed that DRT7_*Ec*_ expression does not affect the growth of the *E. coli* K-12 strain VHB17 (a motile derivative of BW25113^35^) at either 30°C or 37°C (Supplementary Figure S8).

Using VHB17 as the host, we screened the BASEL coliphage collection^36^ and a set of commonly used model phages for sensitivity to the DRT7_*Ec*_ system. An initial screen using overnight cultures to prepare top agar overlays yielded multiple hits, but protection was not reproducible. Re-evaluation of the assay conditions revealed that using exponentially growing cultures for overlays and incubating plates at 30°C gave reproducible results. This optimized screen demonstrated that DRT7_*Ec*_ protects against a broad range of phages (Fig. 5A and B). To further evaluate protection, we monitored growth of VHB17, with and without DRT7_*Ec*_ expression, upon infection with λvir, Bas1, Bas22, or Bas34 at multiplicities of infection (MOIs) of 0, 0.1, 1, and 10 (Fig. 5C). These phages were selected as representative of different taxonomic groups. For λvir, DRT7_*Ec*_ provided near-complete protection at MOIs 1 and 0.1 (Fig. 5C). At MOI 10, protection manifested as slower bacterial growth followed by culture collapse after 7 h, potentially due to reduced expression from the P_*tet*_ promoter in stationary phase or the emergence of λvir escape mutants. For Bas1, Bas22, and Bas34, DRT7_*Ec*_ conferred near-complete protection at MOIs 10, 1, and 0.1 (Fig. 5C), indicating direct defense.

To address the mechanism of DRT7_*Ec*_-mediated defense, we isolated spontaneous escape mutants of λvir and Bas1 that overcame the protective effect of DRT7_*Ec*_ (Fig. 5D). Sequencing of four escape mutants of each phage revealed mutations in genes linked to nucleic acid metabolism. The λvir escape mutants contained nonsense or frameshift mutations in the *gam* gene, which encodes a DNA mimic that inhibits the RecBCD enzyme^37–39^. The Bas1 escape mutants carried missense or frameshift mutations in the Bas01_0050 ORF, which putatively encodes a nucleic acid-targeting phosphoesterase. Although DRT7_*Ec*_ may not be a strict abortive system, the reduced growth rate of DRT7_*Ec*_-expressing cells at high MOI of λvir prompted us to test whether co-expression of DRT7_*Ec*_ and Gam leads to growth inhibition. These assays revealed that co-expression of DRT7_*Ec*_ and Gam indeed reduces growth (Fig. 5E), implying that DRT7_*Ec*_ is activated by Gam. We were unable to perform the analogous experiment with Bas01_0050 as its expression was toxic in *E. coli*.

**Figure 5.**
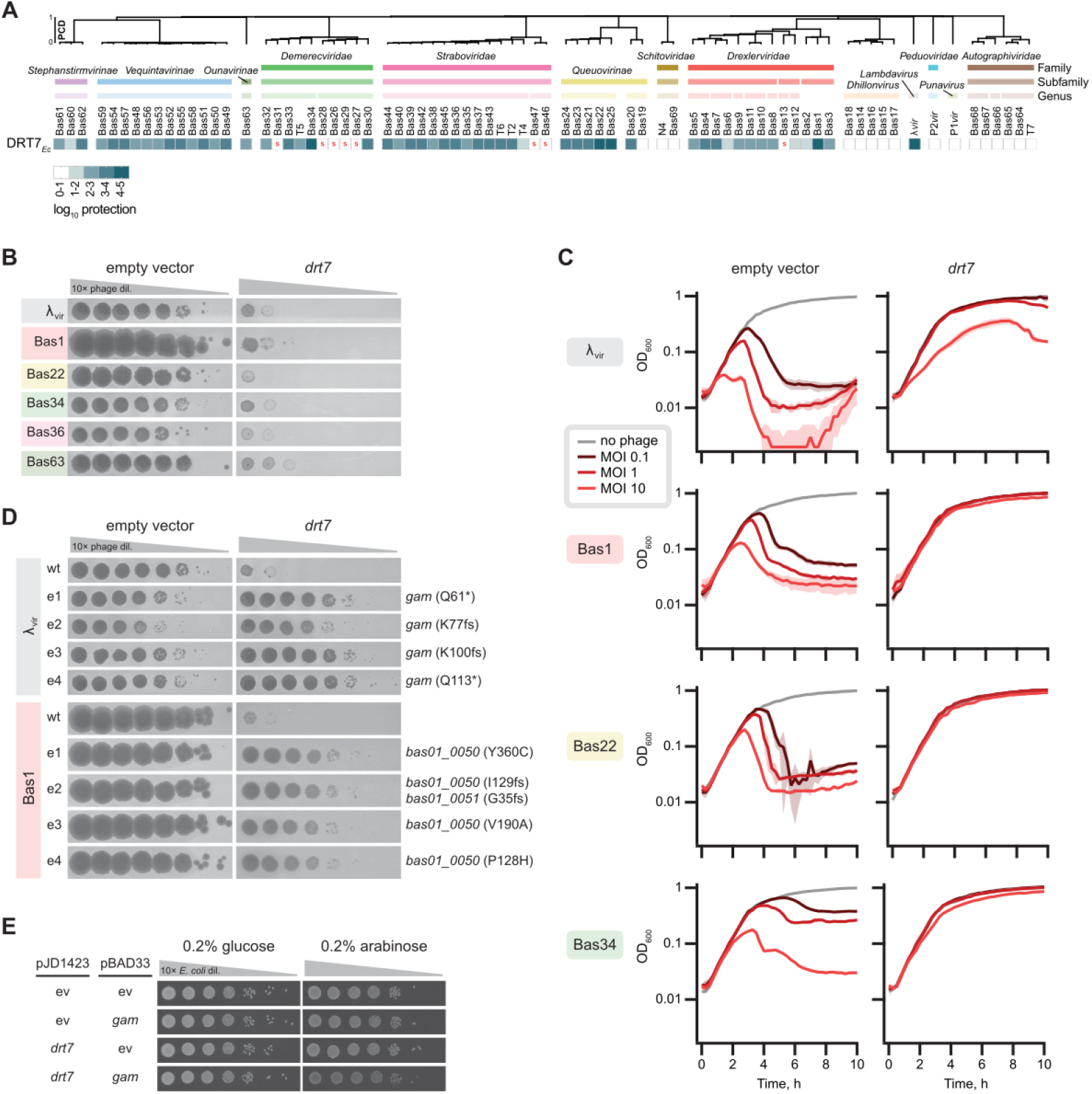
Antiphage activity of DRT7_*Ec*_. (**A**) Defense profile of DRT7_*Ec*_*-*expressing *E. coli*. Top agar lawns of *E. coli* VHB17 harboring the empty pBR322-derived vector pJD1423 or the same plasmid expressing DRT7_*Ec*_ were challenged with 10-fold serial dilutions of indicated phages. The phages that showed sensitivity to the expression of DRT7_*Ec*_ were re-tested using three different transformants. The average EOP value from the three repeats were –log_10_ transformed, generating the log_10_ protection value. A red “s” indicates a reduction in plaque size. The phage order and dendrogram are defined by hierarchical clustering applied to the Proteome Composition Distance matrix^10^. (**B**) Plaque assays on lawns of *E. coli* VHB17 harboring the empty vector or the plasmid containing *drt7*. The indicated phages were 10-fold serially diluted, spotted on top agar plates, and incubated at 30°C. (**C**) Growth of *E. coli* VHB17 harboring the empty vector or the plasmid expressing DRT7_*Ec*_ at 30°C in the absence or presence of indicated phage at MOIs of 0, 0.1, 1, and 10. The curves represent the average of three replicates, using different transformants in each replicate, and the shaded areas indicate the standard deviation. (**D**) Phage mutants escaping the DRT7_*Ec*_ system. Altered genes in the escape phages are indicated to the right. (**E**) Growth of *E. coli* VHB17 expressing DRT7_*Ec*_ and/or Gam.

## DISCUSSION

In this study, we explore the biochemical, structural, and functional features of two bacterial UG10/DRT7 antiphage defense systems. We examined representatives of both forms of this system – the long form (UG10L), where the domains are fused, and the short form (UG10S), where the arch domain is encoded in a separate ORF. A remarkable feature of UG10 enzymes is their ability to produce ssDNA in a template-independent, protein-primed manner and then use it as a template for the synthesis of the second DNA strand. To our knowledge, this is the first enzyme demonstrating such activity. This requires cooperation between two catalytic domains – an RT-like polymerase synthesizing the first DNA strand and an N-terminally fused primase domain producing the complementary strand. Our structural data reveal an inherent flexibility of UG10 proteins, likely crucial for coordinating their polymerase activities.

The RT domain of both tested UG10 systems exhibits high selectivity for deoxythymidine^27^. Nucleotide preference has been previously studied in related Abi/UG RT enzymes, AbiK and AbiA^9,10^. AbiK incorporates all four deoxynucleotides indiscriminately, while AbiA displays a clear bias toward adenosines and cytosines^13^. The mechanism underlying UG10 RT’s preference for dTTP may involve the four highly conserved arginine sidechains that, in both structures, contact the 3′ nucleotide of the DNA strand—the position of the incoming nucleotide. These residues may selectively bind dTTP. Homopolymer synthesis has also been reported for UG28 (DRT9), which synthesizes poly(dA). However, as a Class 2 UG RT, it does so in a template-dependent manner using the associated ncRNA, thus dictating this specificity by the template sequence^16,17^.

To initiate first-strand DNA synthesis, UG10 employs a protein-priming mechanism, similar to other Class 1 Abi/UG RT enzymes studied to date^9,10,28,29^. In these enzymes, priming residues map to flexible loops, which can be located in different subdomains—for example, in the fingers subdomain of AbiK^9^, the palm subdomain of DRT4^28^, and the linker between the RT and αRep domains of AbiA^10^. UG10 differs significantly, positioning its priming residue (Tyr682 in DRT7_*Ec*_ and Tyr691 in UG10S_*Ss*_) within an α-helix of a mobile two-helix module of the αRep domain. We propose that the two-helix module undergoes rearrangement to place the hydroxyl group of the priming tyrosine near the RT active site, enabling the initial step of DNA synthesis.

To our knowledge, UG10 represents the first functionally and structurally characterized antiphage defense system combining the enzymatic activities of an RT-like polymerase and a primase. Primase domains are also found in other defense systems association or C-terminally fused with CRISPR-Cas-associated RTs^6^. Moreover, some members of Class 3 of the Abi/UG RT clade are associated or fused with DnaG-like primase domains, which are also found in certain retron types^7^. An intriguing aspect of UG10’s molecular mechanism is how dsDNA synthesis is orchestrated and how the template strand generated by the RT domain is transferred to the primase for complementary strand synthesis. Despite numerous attempts with several UG10 homologues, we could not obtain high-quality cryo-EM structures of protein-DNA complexes after second-strand synthesis. However, using the available structure of a highly similar primase complexed with DNA products, we modeled the expected trajectory of the ssDNA interacting with the primase domain. The mobility of the priming two-helix module likely positions the DNA covalently linked to it as the template at the primase active site.

While this manuscript was being prepared, another study of the *E. coli* DRT7 system was posted^27^. The major findings of both studies are in agreement—observing protein-primed synthesis of a dsDNA product with a strong nucleotide preference. These works are complementary rather than overlapping. Kim et al. explored, in detail, the catalytic properties of the primase and discovered exonuclease activity associated with the RT domain, endowing the enzyme with proofreading activity and the ability to bypass 8-oxo-dG lesions. Furthermore, their observations—for example, increased primase activity in the absence of the AD—are supported by the insights gleaned from the DRT7_*Ec*_ structure presented here, which clearly demonstrates the blocking of the primase DNA-binding region by one of its domains. However, some of our observations regarding polymerase activity do not fully align with those reported by Kim and colleagues^27^. They found that removal of the AD from DRT7_*Ec*_ abrogated polymerase activity. Conversely, our experiments with UG10S_*Ss*_ (corresponding to this truncation) showed efficient DNA synthesis. These differences may stem from interspecies variations or disparate properties of UG10L and UG10S enzymes.

Notably, our phage infection experiments revealed a broad spectrum of phages against which DRT7 provides defense. The mechanism underlying this defense remains unclear, however, it does not appear to involve triggering cell death, as cultures carrying the system challenged with phages largely maintained growth dynamics similar to uninfected cells. Prior experiments demonstrated that both enzymatic activities of UG10 (DRT7) are required for protection^14^. To gain further insight, we studied phage escape mutants isolated in the presence of active DRT7_*Ec*_ for two phages. Mutations were identified in the *gam* gene of lambda phage and in *bas01_0050* of Bas1. The products of both genes are involved in DNA metabolism. Gam is a known inhibitor of the RecBCD system, which plays a central role in bacterial immunity and DNA recombination/repair. Sensing of Gam-mediated inhibition of RecBCD has been observed with Retron-Ec48^40^. It is possible that the anti-phage activity of UG10 is triggered by defects in RecBCD functionality, but direct interaction between the Gam protein and UG10 is also possible. The product of *bas01_0050* is predicted to be a nucleic acid-targeting phosphoesterase, suggesting that its interplay with UG10 is likely functional rather than direct.

Many questions regarding the UG10 system remain open. The exact role of the AD is unclear. In our DRT7_*Ec*_ structure, it likely prevents or modulates the synthesis of the second strand, requiring rearrangement to enable primase activity on the DNA template made by the RT domain. This suggests that the AD may inhibit the primase domain, keeping it in check until activation is needed. The mechanisms behind UG10’s antiphage defense and how it is triggered still require elucidation. In particular, the role of the poly(dA)/poly(dT) dsDNA product of UG10 in antiphage defense needs to be investigated.

## MATERIALS AND METHODS

### Bacterial strains, plasmids and media

All proteins used in this study were produced in *Escherichia coli* BL21 Star (DE3) cells. Protein-coding sequences – codon-optimized for *Streptococcus salivarius* proteins (UG10S_*Ss*_, WP_104020715.1 and its cognate arch protein WP_104020716.1) and the native sequence from *Escherichia coli* (DRT7_*Ec*_, WP_247156398.1) – were purchased from Invitrogen and BioBasic, respectively. The coding sequences for UG10S_*Ss*_ and DRT7_*Ec*_ were subcloned into pET28 vectors carrying an N-terminal His-tag cleavable by Tobacco etch virus (TEV) protease. Attempts to clone the *S. salivarius* arch protein sequence into pET28 were unsuccessful, likely due to toxicity. Instead, it was subcloned into pGEX-4T1-TEV vector, but the protein could not be purified in soluble, monodisperse form.

All microbiology/phage infection experiments were performed using derivatives of *E. coli* K-12 strain VHB17, a motile derivative of BW25113^41^ that harbors an IS5 element in the intergenic region between *uspC* and *flhDC*^35^. For phage immunity assays, the native sequence encoding DRT7/UG10L (WP_247156398.1) from *E. coli* strain LN97 (DRT7_*Ec*_), along with the 25 upstream nucleotides, was subcloned under the P_*tet*_ promoter in plasmid pJD1423, a derivative of pBR322 in which the *tetA* open reading frame has been replaced by the *rrnB* terminator^34^. A similar construct for the *S. salivarius* UG10S system could not be produced due to difficulties subcloning the arch protein sequence, likely caused by toxicity. The wild-type *gam* and *Bas01_0050* genes were PCR amplified with 20 bp upstream sequence from the genomes of λvir and Bas1, respectively, and cloned by Gibson assembly under the P_*BAD*_ promoter in pBAD33^42^.

*E. coli* strains were grown in liquid or on solid (1.5% wt/vol agar) LB medium (10 g/L tryptone, 5 g/L yeast extract, and 10 g/L NaCl). When required for plasmid selection, the medium was supplemented with 100 μg/ml ampicillin and/or 20 μg/ml chloramphenicol. To repress or induce genes placed under the P_*BAD*_ promoter, the medium was supplemented with 0.2% glucose or 0.2% arabinose, respectively. For top agar overlays, LB agar (0.5% wt/vol agar) supplemented with 20 mM MgSO_4_ and 5 mM CaCl_2_ was used.

### Expression and purification of UG10 proteins

Protein expression was induced overnight with 0.4 mM isopropyl 1-thio-β-D-galactopyranoside (IPTG) at 18°C. Bacterial cells were suspended in 40 mM NaH_2_PO_4_ (pH 7.0), 100 mM NaCl, 5% glycerol, and 10 mM imidazole, supplemented with a mixture of protease inhibitors and viscolase, and incubated on ice with 1 mg/ml lysozyme. After sonication, the cleared lysate was applied to a HisTrap column (Cytiva) equilibrated with 10 mM imidazole, 40 mM NaH_2_PO_4_ (pH 7.0), 500 mM NaCl, and 5% glycerol. Following a wash with 60 mM imidazole, the protein was eluted with 180 mM imidazole. Selected fractions were dialyzed overnight against 20 mM HEPES (pH 7.5), 100 mM NaCl, 5% glycerol, 0.5 mM EDTA, and 1 mM DTT, with or without TEV protease (for UG10S_*Ss*_ and DRT7_*Ec*_, respectively). The sample was then applied to a Heparin column (Cytiva) equilibrated with 20 mM HEPES (pH 7.5), 100 mM NaCl, 5% glycerol, 0.5 mM EDTA, and 1 mM DTT. Bound protein was eluted with a linear gradient of 100-1000 mM NaCl. Pooled, selected fractions were concentrated and applied to a Superdex 200 Increase 10/300 GL gel filtration column (GE Healthcare) equilibrated with 20 mM HEPES (pH 7.5), 100 mM NaCl, 5% glycerol, 0.5 mM EDTA, and 1 mM DTT. Fractions containing pure protein were pooled and concentrated.

### Mass photometry

Mass photometry measurements were performed using a Refeyn TwoMP mass photometer (Refeyn Ltd., Oxford, UK) following the manufacturer’s instructions. Microscope coverslips (No. 1.5H) were cleaned using standard laboratory procedures and assembled into flow chambers immediately prior to measurements. All experiments were conducted at room temperature. The instrument was calibrated using a protein mass standard in the same buffer as the samples, prior to data acquisition.

Samples were diluted to nanomolar concentrations in measurement buffer containing 20 mM HEPES (pH 7.5), 100 mM NaCl, 1 mM EDTA, and 1 mM DTT. A buffer blank was recorded to establish the background signal, followed by introduction of diluted samples into the flow chamber. Movies were acquired for 60–120 s using standard settings. Data analysis—including particle landing event detection and conversion into mass distributions based on the calibration curve—was performed using AcquireMP and DiscoverMP software (Refeyn Ltd.). Reported masses represent the mean values obtained from replicate measurements.

### Cryo-EM sample preparation and data collection

UG10S_*Ss*_ (3 μl, 3 mg/ml) was applied to a glow-discharged Quantifoil R2/1 mesh 200 Au grid and vitrified in liquid ethane using an FEI Vitrobot Mark IV (Thermo Fisher Scientific) at 4°C with 95% humidity and a 4 s blot time. Initial grid screening was performed using a Glacios electron microscope (Thermo Fisher Scientific) operating at 200 kV (Center of New Technologies, University of Warsaw). Subsequent data collection was performed with a Titan Krios G3i electron microscope (Thermo Fisher Scientific) operating at 300 kV, equipped with a BioQuantum energy filter (20 eV energy slit) and a K3 camera (Gatan), at the SOLARIS National Synchrotron Radiation Centre (Krakow, Poland). A total of 5307 movies were recorded in counting mode with a physical pixel size of 0.84 Å (nominal magnification of 105,000×), 70 μm objective aperture, and a defocus range of −2.1 μm to −0.6 μm (0.3 μm steps). The total dose, fractionated into 40 frames, was 40.87 e/Å^2^, with a dose per frame of 1.02 e/Å^2^.

UG10S_*Ss*_ D631N (3 μl, 3 mg/ml) was vitrified, screened, and imaged using the same conditions as the wild-type protein. A total of 5860 movies were recorded with 20° stage tilt in counting mode with a physical pixel size of 0.84 Å (nominal magnification of 105,000×), 70 μm objective aperture, and a defocus range of −2.0 μm to −1.4 μm (0.2 μm steps). The total dose (fractionated into 40 frames) was 41.56 e/Å^2^, and the dose per frame was 1.04 e/Å^2^.

DRT7_*Ec*_ (3 μl, 4 mg/ml) was applied to a glow-discharged Quantifoil R2/1 mesh 200 Cu grid and vitrified using the same conditions described above. The grid was imaged with a Glacios 2 electron microscope (Thermo Fisher Scientific) operating at 200 kV and equipped with a Falcon 4i camera at the International Institute of Molecular and Cell Biology in Warsaw. A total of 8097 movies were recorded in counting mode with a physical pixel size of 0.72 Å (nominal magnification of 190,000×), 20 μm objective aperture, and a defocus range of −1.9 μm to −1.0 μm (0.3 μm steps). The total dose (fractionated into 891 fractions) was 31.16 e/Å^2^.

### Cryo-EM data processing and interpretation

Cryo-EM images were processed with RELION-5.0.0^43^ and cryoSPARC 4.7.1^44,45^. For all datasets, raw movies were motion-corrected and binned 2× using RELION’s implementation of MotionCor2 (Supplementary Figs. 2, 3, and 7). For the dataset collected using Glacios 2, 27 dose fractions were grouped into one, resulting in 33 frames with a dose per frame of 0.94 e/Å^2^. Contrast transfer function (CTF) was fitted using CTFFIND4.1^46^ for the UG10S_*Ss*_ and DRT7_*Ec*_ datasets.

For the UG10S_*Ss*_ dataset, 2,331,461 particles were selected with crYOLO^47^ and binned to 3.36 Å/pix, then subjected to three rounds of reference-free 2D classification in cryoSPARC (Supplementary Fig. 2). From these, 1,640,495 particles were re-extracted in RELION with a pixel size of 1.68 Å/pix and subjected to four rounds of heterogeneous refinement in cryoSPARC using five volumes generated with the *ab initio* procedure. Non-uniform refinement of particles from the best class yielded a 4.22 Å reconstruction. Bayesian polishing (with re-extraction at an unbinned pixel size of 0.84 Å/pix) in RELION, followed by non-uniform refinement with global and local CTF refinements in cryoSPARC, improved resolution to 3.73 Å. Two additional heterogeneous refinement rounds in cryoSPARC, using one high-quality volume and four decoy volumes, allowed the selection of 235,883 particles that produced a 3.69 Å map, which improved to 3.56 Å after a second round of Bayesian polishing in RELION. 3D variability analysis in cryoSPARC revealed two major volume classes – one with well-defined density corresponding to the two-helix module in the αRep domain, and one in which this module is poorly resolved, suggesting conformational flexibility. The 103,501 particles from the former class were used to produce the final reconstruction with a resolution of 3.68 Å. The map was globally sharpened with a B-factor of –196. Individual domains of the UG10S_*Ss*_ structure, predicted using AlphaFold 2^48^, were fitted into the map and the resulting model was further manually edited in Coot v0.9.8.1^49^ and refined using real space refinement in Phenix^50^. The entire protein chain is visible except for residues 237-247. A single-stranded DNA fragment of 9 pyrimidine nucleotides could be traced in the map and was built as poly(dT) DNA, consistent with the observed nucleotide preference of UG10S_*Ss*_. For the DRT7_*Ec*_ dataset, 2,006,829 particles were selected with crYOLO, binned to 2.87 Å/pix, and subjected to two rounds of reference-free 2D classification in cryoSPARC (Supplementary Fig. 3).

A total of 630,428 selected particles were re-extracted in RELION with a pixel size of 1.43 Å/pix and subjected to two more rounds of 2D classification. Non-uniform refinement of 513,337 selected particles yielded a 3.40 Å reconstruction. After Bayesian polishing in RELION (re-extraction with an unbinned pixel size of 0.72 Å/pix), polished particles were imported into cryoSPARC. Non-uniform refinement with global and local CTF refinements improved the resolution to 2.86 Å. After two rounds of heterogeneous refinement using one high-quality and two decoy volumes, a subset of 293,305 particles was selected, producing a 2.84 Å map. This subset was pruned through 3D classification to remove particles with poorly resolved domains, yielding a 2.91 Å map that improved to 2.85 Å after a second round of Bayesian polishing in RELION. The map was globally sharpened with a B-factor of –105. To solve the structure, a model generated using AlphaFold3 was docked in the map, manually adjusted in Coot, and refined using real space refinement in Phenix. The fragment encompassing residues 1528-1680 could not be traced, as corresponding density was absent. A single-stranded DNA fragment of 10 nucleotides could be visualized and was built as poly(dT) DNA.

For the UG10S_*Ss*_ D631N dataset, CTF was fitted using the patch CTF tool in cryoSPARC (Supplementary Fig. 7). Template-based particle picking was performed using 2D classes from initial processing as templates. A total of 4,974,821 particles were extracted with a pixel size of 3.36 Å/pix and subjected to two rounds of reference-free 2D classification, followed by three rounds of heterogeneous refinement using five volumes generated *ab initio*. The selected subset of 1,287,064 particles was re-extracted with a pixel size of 1.68 Å/pix and subjected to two more rounds of heterogeneous refinement, leading to a subset of 638,484 particles. Non-uniform refinement yielded a 3.66 Å reconstruction. Bayesian polishing (with re-extraction at an unbinned pixel size of 0.84 Å/pix) in RELION, followed by non-uniform refinement with global and local CTF refinements in cryoSPARC, improved resolution to 3.42 Å. Two additional rounds of heterogeneous refinement using the same volumes as previously yielded a subset of 483,104 particles, producing a 3.40 Å map. These particles were subjected to 3D classification in cryoSPARC. The four resulting classes were similar in size and overall quality, but differed in local map quality around the protein-priming two-helix module and the αRep domain. Further analysis was performed using 125,613 particles from the class with the best-resolved features, yielding a 3.62 Å reconstruction. Particles were imported into RELION for another round of Bayesian polishing, and the resulting polished particles were imported into cryoSPARC for a final round of non-uniform refinement, yielding a 3.52 Å reconstruction, globally sharpened with a B-factor of –158. For structure determination, the wild-type UG10S_*Ss*_ structure was used as a starting model, manually adjusted in Coot, and refined using real space refinement in Phenix. The map allowed visualization of the protein spanning residues 1-695, except for several short loops. A fragment of the αRep domain encompassing residues 744-853 could be docked into the map by rigid body fitting of a corresponding fragment from the UG10S_*Ss*_ structure.

All reported resolutions were estimated from gold-standard masked Fourier Shell Correlations (FSC) at the 0.143 threshold. Structures and maps were rendered in UCSF ChimeraX 1.10^51^ and PyMOL (PyMOL Molecular Graphics System, Version 2.5.4 Schrödinger, LLC).

### Polymerase nucleotide preference and activity assay

To determine the nucleotide preference of UG10S_*Ss*_ polymerases, proteins (1 μM) were mixed with individual deoxynucleotides (200 μM) in a reaction buffer containing 50 mM Tris (pH 8.5), 100 mM NaCl, 5 mM DTT, and 3 mM MgCl_2_. Reactions were incubated at 37°C for 1 h and stopped by adding 40 mM EDTA. Samples were then treated with proteinase K to remove the enzyme. All reactions were performed in triplicate. Hydrolysis products were analyzed by 10% denaturing Tris-borate EDTA (TBE)-urea polyacrylamide gel electrophoresis and visualized with SYBR Gold staining, followed by fluorescence detection using an Amersham Typhoon biomolecular imager (Cytiva). The same method was used to assess the polymerase activity of UG10S_*Ss*_ point-substituted variants with single 15-minute time point.

To monitor DNA polymerization progression over time, wild-type proteins were first incubated in the reaction buffer (as described above) with 0.5 μM fluorescein-dUTP at 37°C for 15 minutes to label co-purifying DNA. The reaction was then supplemented with 200 μM dTTP and incubated at 37°C for 7.5, 15, 30, 60, or 120 minutes. Reactions were stopped by adding 40 mM EDTA and treating with proteinase K. Hydrolysis products were analyzed by 10% denaturing TBE-urea polyacrylamide gel electrophoresis and visualized by fluorescence detection with the Amersham Typhoon imager. A DNA size marker was generated by digesting ΦX174 DNA (New England BioLabs, N3023S) with HaeIII restriction enzyme, yielding fragments of 72-1353 nucleotides.

### Analysis of nucleic acid products of UG10 reverse transcriptases

UG10S_*Ss*_ protein or its primase-deficient variant (D72A/D74A, AXA) (1 μM) was first incubated at 37°C for 15 minutes in the reaction buffer (as described above) with 0.5 μM fluorescein-dUTP. Following this, 200 μM dTTP in reaction buffer was added, and the mixture was incubated at 37°C for 1 hour. Next, dATP or dCTP (50 μM final concentration) or buffer alone was added, and incubation continued for another hour. The reaction was stopped by adding 40 mM EDTA and treating with proteinase K. Nucleic acid products were then ethanol-precipitated and resuspended in water. To determine their composition, DNA products were treated with nucleases. For ssDNA digestion, 500 ng of purified DNA was mixed with 10 U of S1 nuclease in a buffer containing 40 mM sodium acetate pH 4.5, 300 mM NaCl, 2 mM ZnSO_4_ and incubated for 30 minutes at room temperature. The reaction was stopped with 3 μl of 500 mM EDTA and further incubated at 70°C for 10 minutes. To cleave dsDNA, 250 ng of purified DNA was incubated with 1 μl of dsDNase in the buffer supplied by the manufacturer for 2 minutes at 37°C. The reaction was stopped with 1 μl of 100 mM DTT followed by incubation at 55°C for 5 minutes. The products were analyzed by 10% denaturing TBE-urea polyacrylamide gel electrophoresis and visualized by fluorescence detection using the Amersham Typhoon imager.

### Phage infection assays

To identify phages sensitive to the DRT7_*Ec*_ defense system, *E. coli* VHB17 (BW25113 *uspC-flhDC*::IS5) carrying either the empty pJD1423 vector^34^ or the same plasmid expressing DRT7_*Ec*_ from the constitutive P_*tet*_ promoter were grown overnight at 37°C in LB medium containing 100 μg/ml ampicillin. Cultures were diluted to OD_600_ = 0.03 in the same medium and grown at 37°C until OD_600_ ≈ 0.7. Cells were mixed with top agar (final concentration 0.075 OD600 units/ml) and overlaid onto 12 × 12 cm LB agar plates. Individual phage stocks were 10-fold serially diluted in SM buffer (0.05 M Tris-HCl pH 7.5, 0.1 M NaCl, 0.01 M MgSO_4_), and 2.5 μL of each dilution was spotted onto the top agar. Spots were air-dried for 30 min at room temperature, and plaque formation was scored after 16 h of incubation at 30°C. Sensitive phages were re-tested in triplicate using different transformants for each repeat. Plaques were counted, and the efficiency of plating (EOP) was calculated for each repeat by dividing plaque-forming units for the DRT7_*Ec*_-expressing system by those for the empty vector. Average EOP values from the three repeats were –log10-transformed and used to generate the heatmap (log10 protection).

For monitoring phage infection of liquid cultures, three different transformants of VHB17 carrying either the empty vector or the DRT7_*Ec*_-expressing plasmid were grown overnight at 37°C in LB medium containing 100 μg/ml ampicillin, 10 mM MgSO_4_, and 2.5 mM CaCl_2_. Cultures were diluted to OD_600_ = 0.03 and grown at 30°C to OD_600_ ≈ 0.5 (Ultrospec 7000 spectrophotometer, Biochrom). Cells were then diluted to OD_600_ = 0.075, and 100 μL was added to wells of a 96-well plate containing either 10 μL of SM buffer or 10 μL of appropriate phage dilution. Phage dilutions (prepared in SM buffer from freshly prepared stocks) were used to achieve multiplicities of infection (MOI) of 0.1, 1, and 10. Bacterial growth was monitored at 30°C in a SPECTROstar Nano plate reader (BMG LABTECH) with double-orbital shaking at 400 rpm, with OD600 readings taken every 15 min for 10 h.

### Isolation and genome sequencing of phage escape mutants

To identify mutant phages escaping DRT7_*Ec*_-mediated defense, we purified individual wild-type Bas1 and λ_vir_ clones from our laboratory stocks using VHB17 as the host through three consecutive rounds of single-plaque streaks on top-agar overlays. Following preparation of phage stocks^35^, potential escape mutants were identified as discrete plaques appearing against a background of poor plaque formation in serial dilution plaque assays using top-agar overlays prepared with exponentially growing DRT7Ec-expressing VHB17 cells. Candidate escape mutants were purified by three consecutive rounds of single-plaque streaks and then used to prepare phage stocks. Escape was confirmed by serial dilution plaque assays, and genomic DNA was isolated from the mutants and corresponding wild-type clone using the Phage DNA Isolation Kit (Norgen Biotek). Phage genome sequencing was performed by Seqvision (Vilnius, Lithuania) using Oxford Nanopore technology. Briefly, genomic DNA was end-repaired with 3’ dA addition and 5’ phosphorylation. Adapters were ligated using a T/A ligation-based method without fragmentation, and the genome was amplified using barcoded primers annealing to the adapters. Samples were purified, pooled, and nanopore native adapters were ligated (kit 14 chemistry). Sequencing was performed on R10.4.1 nanopore, and basecalling used the super-accuracy v5.2.0 model on Dorado v1.0.0. Adapters were removed and reads demultiplexed using Dorado v1.0.0 (https://github.com/nanoporetech/dorado) with default parameters. After removing the lowest 10% quality reads using filtlong v0.2.1 (https://github.com/rrwick/Filtlonggithub.com/rrwick/Filtlong), reads were subsampled to four datasets of 100-200x coverage using Autocycler subsample v0.4.0 (https://github.com/rrwick/Autocycler) and assembled using flye v2.9.4 (https://github.com/mikolmogorov/Flye) with default parameters. Subassemblies were used to create a consensus assembly using Autocycler v0.4.0, and the assembly was polished using medaka v2.1.1 with default parameters (https://github.com/nanoporetech/medaka). Mutations in phages relative to the reference Bas1 and λ genomes (MZ501051 and NC_001416) were identified using breseq v0.39.0^52^ with default parameters.

### Bacterial growth assays

To investigate the influence of DRT7_*Ec*_ expression on VHB17 growth, three different transformants carrying either the empty pJD1423 vector or the DRT7_*Ec*_-expressing plasmid were grown overnight at 37°C in LB medium containing 100 μg/ml ampicillin. Cultures were diluted to OD_600_ = 0.03 in 5 ml of the same medium and grown at either 30°C or 37°C until OD_600_ = 0.5 (Ultrospec 7000 spectrophotometer, Biochrom). Cultures were back-diluted to OD_600_ = 0.03, and 100 μL were transferred to a well in a 96-well plate. Growth was monitored at either 30°C or 37°C for 8 h in a Synergy H1 (BioTek) plate reader with double-orbital shaking at 425 rpm, measuring OD_600_ every 15 min.

To investigate the effect of co-expression of DRT7_*Ec*_ and Gam, VHB17 was transformed with combinations of the empty pJD1423 and pBAD33 vectors and their DRT7 and *gam*-containing derivatives. Single colonies were inoculated into LB supplemented with 20 μg/ml chloramphenicol, 100 μg/ml ampicillin, and 0.2% glucose and grown overnight at 37°C. Cultures were diluted to OD_600_ = 0.03 in 5 ml of LB medium containing 20 μg/ml chloramphenicol, 100 μg/ml ampicillin, and grown at 30°C until OD_600_ ≈ 0.5. Cultures and 10-fold serial dilutions were spotted (5 μL) onto LB agar plates containing 20 μg/ml chloramphenicol, 100 μg/ml ampicillin, and either 0.2% glucose or 0.2% arabinose. Plates were imaged after 20 h of incubation at 30°C.

## AUTHOR CONTRIBUTIONS

MF solved and analyzed structures, supervised the project, and co-wrote the manuscript. VKV purified proteins, prepared cryo-EM samples, and performed biochemical experiments. MCC processed the cryo-EM data. MŠ and JR purified proteins. AE performed bioinformatic analyses. VH analyzed the data and co-wrote the manuscript. MJOJ performed antiphage activity experiments, analyzed the data, co-wrote the manuscript, and supervised the project. MN analyzed structures, co-wrote the manuscript, and supervised the project.

## ACKNOWLEDGMENTS

This work was financed by the statutory funding of the International Institute of Molecular and Cell Biology (IIMCB). IIMCB core facilities (IN-MOL-CELL infrastructure) were funded by the European Union and co-financed under the European Funds for Smart Economy 2021-2027 (FENG). This publication was supported by the Polish Ministry of Education and Science project, “Support for research and development with the use of research infrastructure of the National Synchrotron Radiation Centre SOLARIS” (contract no. 1/SOL/2021/2). We thank the SOLARIS Centre for access to the cryo-EM Beamline and Tomasz Góral (CeNT, University of Warsaw) for access to the Glacios electron microscope used for cryo-EM grid screening.

The computations were enabled by the Berzelius resource (Knut and Alice Wallenberg Foundation at the National Supercomputer Centre) and by resources provided by the National Academic Infrastructure for Supercomputing in Sweden (NAISS) (Swedish Research Council grant agreement no. 2022-06725). This work was further supported by the Knut and Alice Wallenberg Foundation (project grant 2020-0037 to V.H.), the Swedish Research Council (Vetenskapsrådet) grants (2021-01146 and 2024-06059 to V.H.), the Göran Gustafsson Foundation for Research in Natural Sciences and Medicine (the Göran Gustafsson Prize to V.H.), and the Carl Trygger Foundation (CTS24:3450 to M.J.O.J.).

