## Supplementary Text and Figures for "Structures and enzymatic mechanisms of DRT7/UG10 antiphage reverse transcriptases"

### SUPPLEMENTARY INFORMATION

### SUPPLEMENTARY TEXT

#### Relative positions of domains in UG10S$_{Ss}$ and DRT7$_{Ec}$

Within the enzymatic cores of UG10/DRT7 proteins, the PriL-CTD domain is connected to the primase and RT domains by flexible, unstructured linkers, allowing for variable domain arrangements. Different arrangements were observed in the structures of UG10S$_{Ss}$ and DRT7$_{Ec}$.

In the UG10S$_{Ss}$ structure, the PriL-CTD is positioned near the interface between the palm and fingers subdomains of RT, while the neighboring primase domain is close to the palm subdomain (Supplementary Fig. 5A). This arrangement creates two interfaces between the RT domain and primase/PriL-CTD. The first, a small interface, involves the second α-helix and a preceding loop of the primase (residues 37 and 50) and the tip of a helix from the palm subdomain (residues 519 and 521). The second interface comprises the second to last helix and following loop of the PriL-CTD domain (residues 327-342), which interact with a region encompassing both palm and fingers subdomains (residues 386, 389, 471, 475, 483, 484, 491, 493, 494). Furthermore, the primase domain is positioned near the two-helical module of the αRep domain (Supplementary Fig. 5B).

In contrast, in DRT7$_{Ec}$, the domains are arranged in a roughly square plan, with the PriL-CTD positioned opposite the fingers subdomain and the primase opposite the palm subdomain (Supplementary Fig. 5A). The interface between the primase and RT domain involves the edge of a β-sheet in the N-terminal part of the primase, which is in proximity to the termini of two helices from the palm subdomain (residues 516, 613, and 614). Additional contacts include an interaction between Asp164 and Tyr499, and two contacts formed by the linker between primase and PriL-CTD: Arg242 stacking against Tyr469, and Arg245 interacting with Asp384. This arrangement positions the primase domain further from the αRep domain, resulting in a more open conformation of this region (Supplementary Fig. 5B), and substantially increasing the distance between the primase and the two-helical module of the αRep.

#### The arch domain of DRT7$_{Ec}$

The structure of DRT7$_{Ec}$ reveals a 941-amino acid C-terminal accessory domain (AD) forming a large arch over the UG10S-like core (Supplementary Fig. 6A). The portion of the AD visible in the cryo-EM maps is predominantly α-helical, comprising 39 helices, along with several small β-sheets: a four-stranded β-sheet (residues 1063-1120), a β-hairpin (residues 1185-1199), and two adjacent antiparallel β-sheets of three or five strands (residues 1370-1435).

Five globular modules can be identified within the AD (Supplementary Fig. 6A,B). The first (residues 837-990) is an N-terminal portion with a helical arrangement similar to the αRep domain, representing a continuation of that domain's solenoid structure. This N-terminal subdomain interacts with the second module (residues 991-1271), which is less compact and contains several β-strand elements. This module contacts the PriL-CTD domain via residue pairs Pro1103-Tyr322 and Asn1154-Leu284/Arg289 (Supplementary Fig. 6C). The third module (residues 1272-1468), composed of α-helices and two β-sheets, is the only AD element interacting with multiple subdomains of the primase-RT core (residue pairs: Thr1274-Ser159,

Arg1411-Glu513, Glu1420-Thr695) (Supplementary Fig. 6E,F). Additionally, residues 1355-1362 are in close proximity to Lys73 of the primase domain. The fourth visible module (residues 1469-1771) forms a protrusion from the overall $DRT7_{Ec}$ globular structure, with the C-terminal region contacting the primase domain (Val1723-Thr137, Phe1724-Phe232, Lys1726-Gly230) (Supplementary Fig. 6D). The region spanning residues 1528-1681 could not be modeled, but AlphaFold3 predictions suggest it forms a compact domain projecting from module 4 via flexible loops (Supplementary Fig. 6B).

The AD interacts not only with individual primase-RT domains but also appears to influence their relative positions, as evidenced by the differences in domain arrangement between $UG10S_{Ss}$ and $DRT7_{Ec}$ (Supplementary Fig. 5). Insertion of AD module 3 between the primase domain and the two-helix module of αRep likely inhibits or obstructs cooperation between these enzymatic activities (Supplementary Fig. 6E,F) and may underlie the observed structural differences. Interestingly, the AlphaFold3 model of the $UG10S_{Ss}$ complex with its cognate arch protein (AP) shows a different arrangement, where module 3 of the AD is not inserted between these domains.

Structural differences between the enzymatic modules in $DRT7_{Ec}$ and $UG10S_{Ss}$ also influence protein-DNA interactions, particularly at the 5' end of the strand. In the $UG10S_{Ss}$ structure, the first two nucleotides interact with residues Phe62, Phe168, and Ser61 of the primase subdomain (Fig. 3E). Residues Arg287 and Asn320 of PriL-CTD also contact the base and sugar ring of nucleotide 4. Poor density between nucleotides 2 and 4, and an inability to model the intervening DNA fragment, suggest flexibility and potential heterogeneity in the length of DNA strands co-purifying with the protein. In contrast, the initial nucleotides of the $DRT7_{Ec}$ DNA product form an extensive network of contacts with residues in both the palm subdomain and the AD (Fig. 3E).

### SUPPLEMENTARY FIGURES

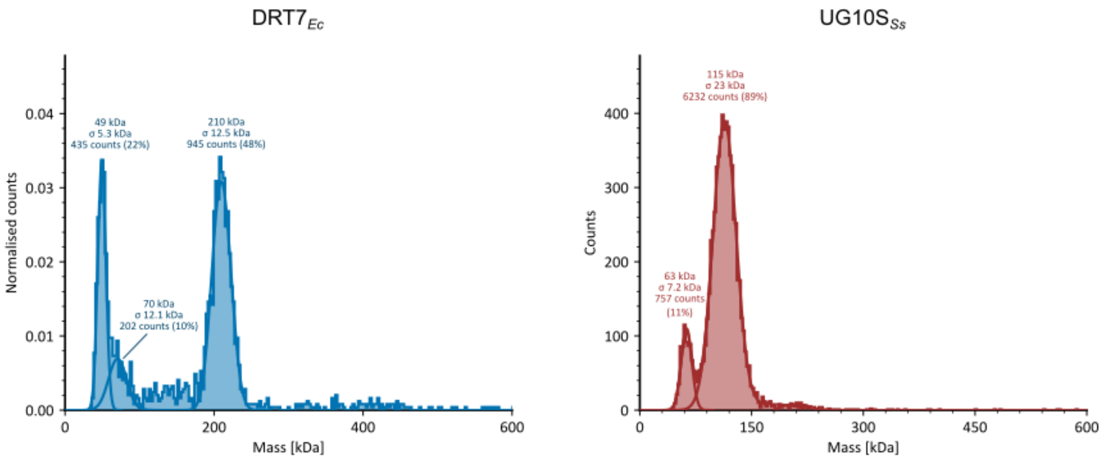


**Supplementary Figure 1. Mass photometry**. Molecular mass distribution histograms of the DRT7$_{Ec}$ and UG10S$_{Ss}$ samples.

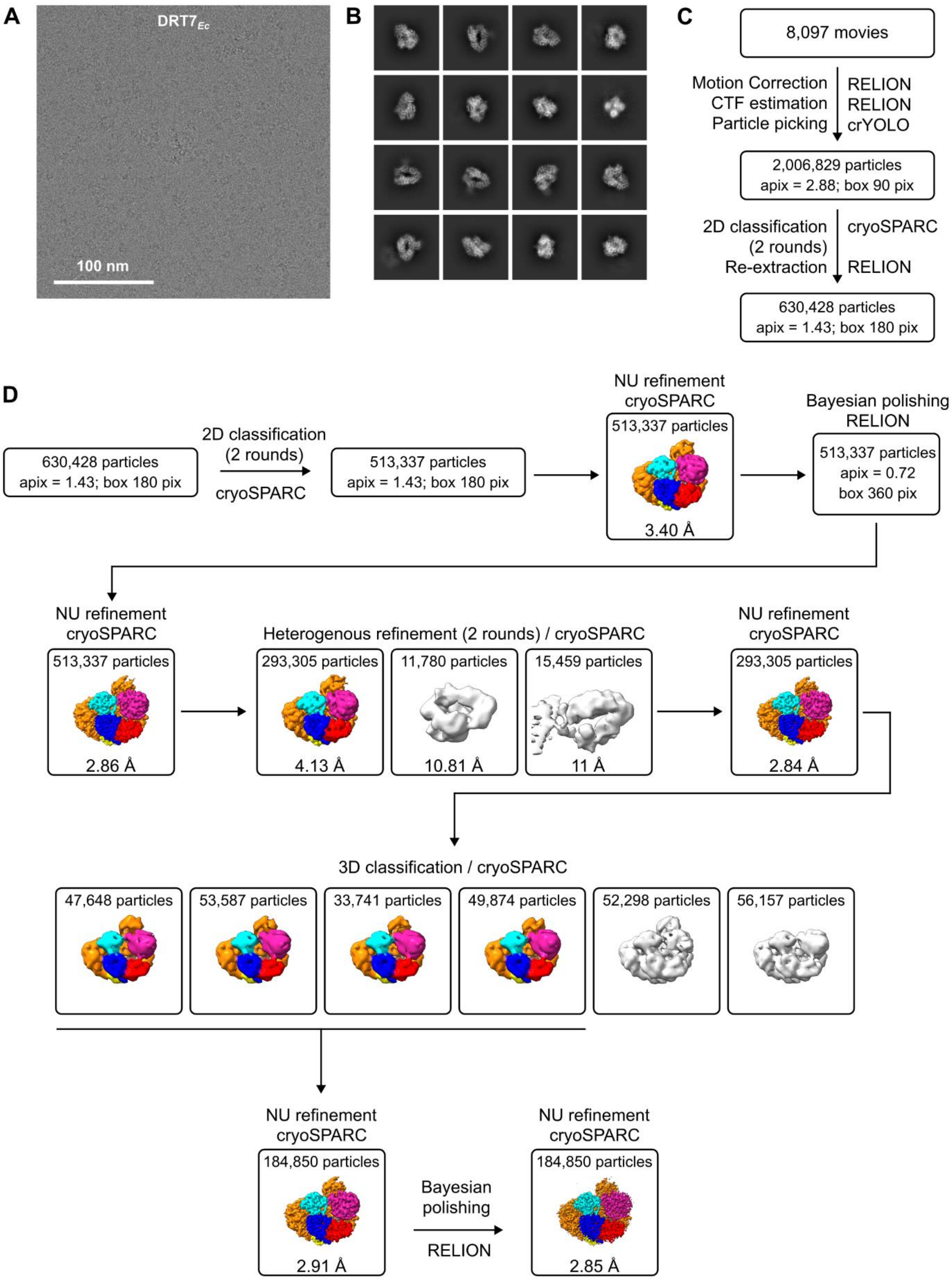


**Supplementary Figure 2. Cryo-EM data processing for DRT7$_{Ec}$.** (**A**) Representative cryo-EM micrograph. (**B**) Representative 2D class averages. (**C**) Initial processing steps (preprocessing, particle picking, and curation). (**D**) Three-dimensional reconstruction pipeline with intermediate maps and 3D classes color-coded as in Fig. 1. Statistics from the final round of heterogeneous refinement are provided.

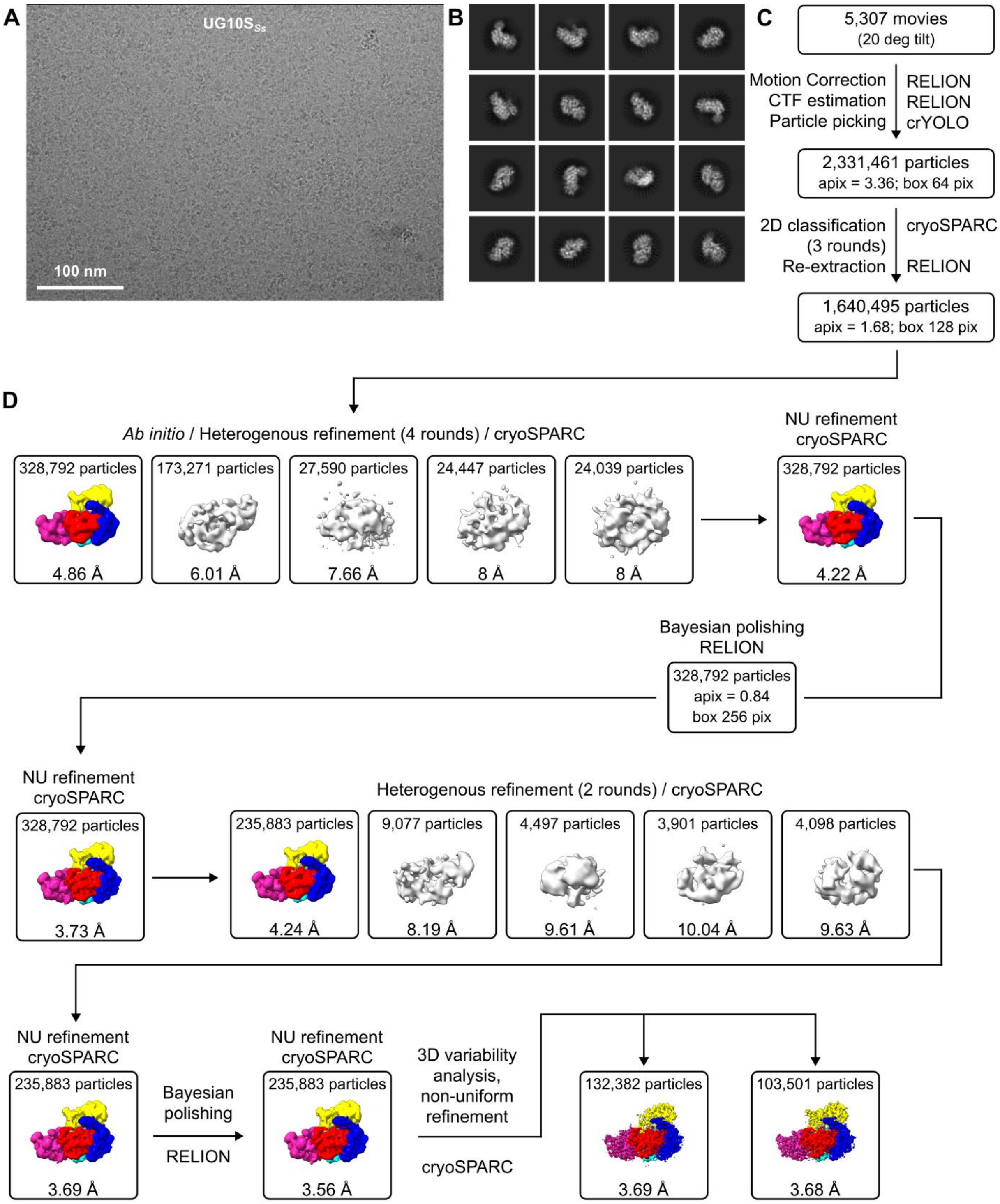


**Supplementary Figure 3. Cryo-EM data processing for UG10S$_{Ss}$.** (**A**) Representative cryo-EM micrograph. (**B**) Representative 2D class averages. (**C**) Initial processing steps (preprocessing, particle picking, and curation). (**D**) Three-dimensional reconstruction pipeline with intermediate maps and 3D classes color-coded as in Fig. 1. Statistics from the final round of heterogeneous refinement are provided.

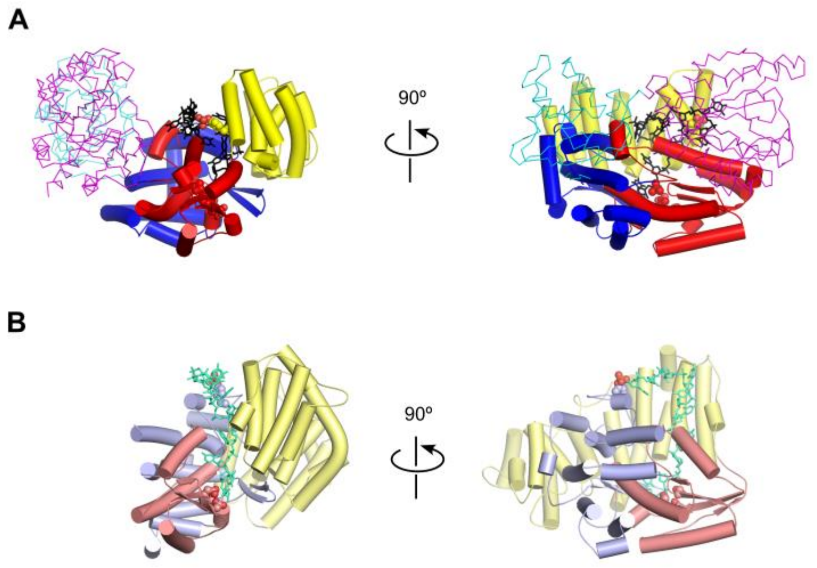


**Supplementary Figure 4. Comparison of DRT7$_{Ec}$ (A) and *Ll*-AbiK (B).** Structures of DRT7$_{Ec}$ (this work) and *Ll*-AbiK (PDB ID: 7R07) were superimposed. RT and αRep domains of DRT7$_{Ec}$ are shown in cartoon representation (colored as in Fig. 1), with lighter shades of the same colors for *Ll*-AbiK. DNA strands are shown as sticks in black (DRT7$_{Ec}$) and teal (*Ll*-AbiK). Primase and PriL-CTD domains are shown in wire representation. The AD is omitted from (A) for clarity.

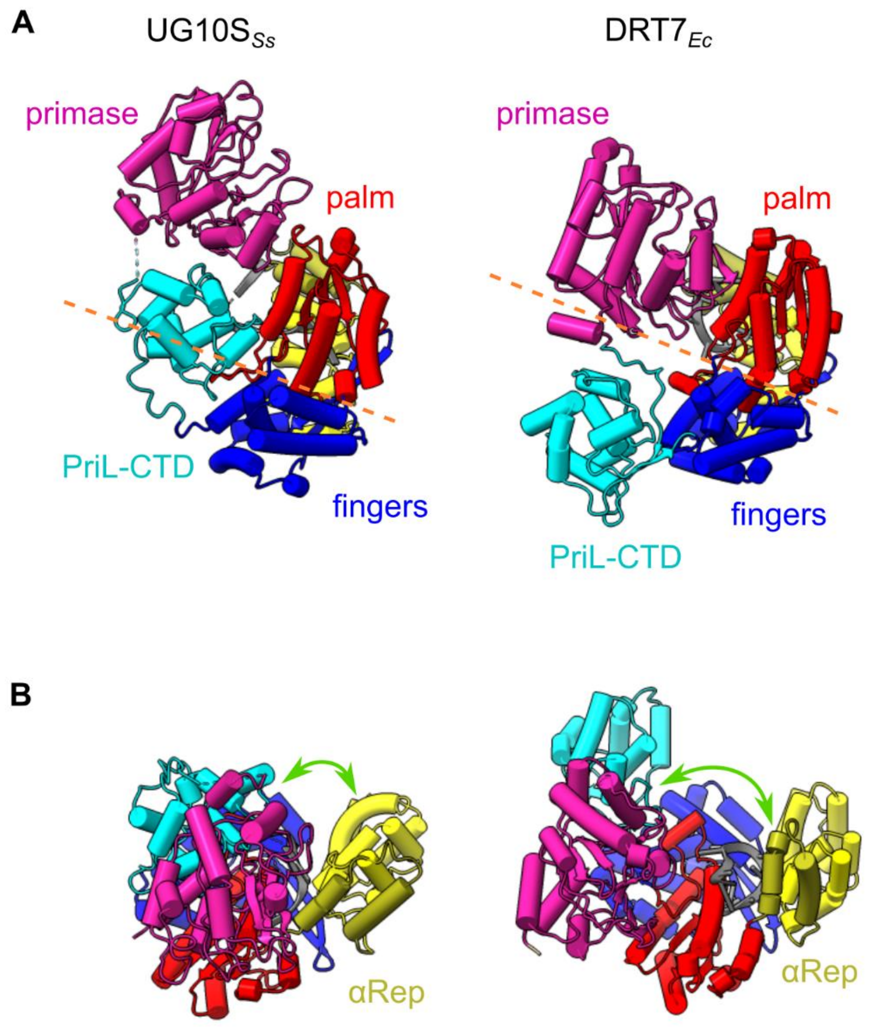


**Supplementary Figure 5. Domain arrangements in UG10S$_{Ss}$ and DRT7$_{Ec}$.** (**A**) Arrangement of primase, PriL-CTD, and RT domains in cryo-EM structures. The plane between the palm and fingers subdomains of RT is indicated by a dashed orange line. (**B**) Relative arrangement of primase, PriL-CTD, and αRep domains, with distances between αRep and other domains indicated by green arrows.

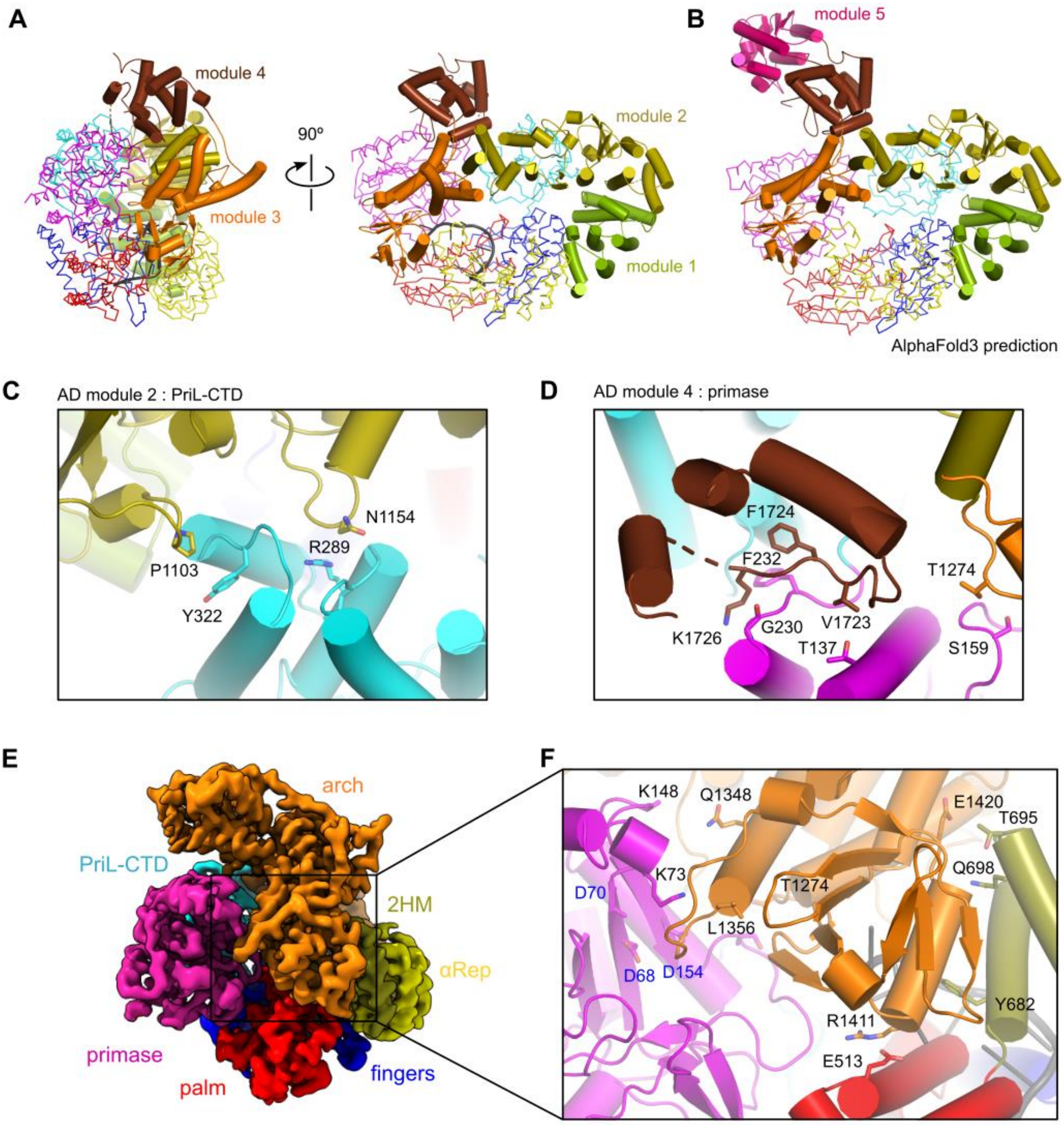


**Supplementary Figure 6. Structure of the Arch Domain (AD) of DRT7$_{Ec}$.** (**A**) AD depicted in the context of the full protein, with AD subdomains shown in different colors. The primase-RT-αRep portion is shown in wire representation (colored as in Fig. 1). (**B**) AlphaFold3 prediction of DRT7$_{Ec}$ structure, with unmodeled regions shown in pink. (**C, D**) Interfaces between modules two (C) and five (D) of the AD and the enzymatic domains. Interacting residues are shown as sticks and labeled. (**E**) Cryo-EM reconstruction of DRT7$_{Ec}$ visualizing the AD subdomain (residues 1272-1468) inserted between the primase and αRep domains. (**F**) Close-up of the interface between the 1272-1468 AD subdomain and the enzymatic domains of DRT7$_{Ec}$. Interacting residues are shown as sticks and labeled, with primase active site residues labeled in blue.

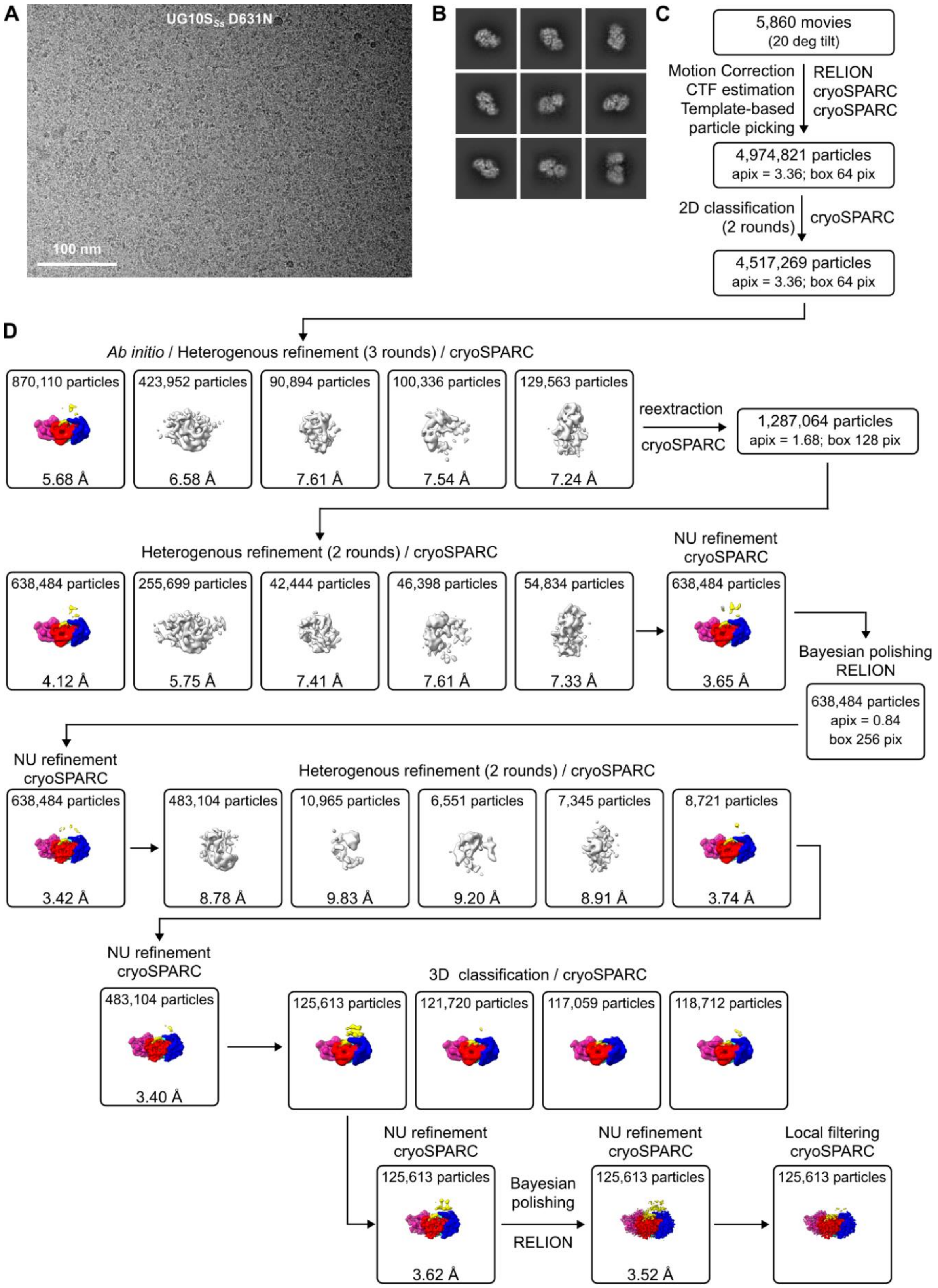


**Supplementary Figure 7. Cryo-EM Data Processing for UG10SSs D631N.** (**A**) Representative cryo-EM micrograph. (**B**) Representative 2D class averages. (**C**) Initial processing steps (preprocessing, particle picking, and curation). (**D**) Three-dimensional reconstruction pipeline with intermediate maps and 3D classes color-coded as in Fig. 1. Statistics from the final round of heterogeneous refinement are provided.

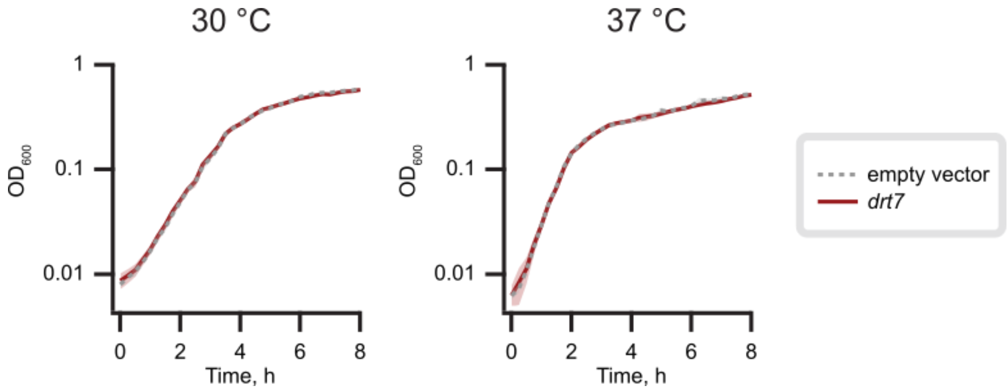


**Supplementary Figure 8. Liquid growth assays of the DRT7*$_{Ec}$*-expressing *E. coli* K-12 strain VHB17.**
